# Reversing aging-like 3D genome disorganization in a *Drosophila* interphase model

**DOI:** 10.64898/2026.07.16.739007

**Authors:** Alexey V. Onufriev, Junkai Zhang, Igor V. Sharakhov, Igor S. Tolokh

## Abstract

Recent experimental evidence suggests that aging may arise from the progressive deterioration of the epigenetic landscape, while reversing the trend can result in cell and tissue rejuvenation. A mechanistic understanding of how restoration of a key component of this landscape – the 3D structure of the genome – can be accomplished is lacking. Here we investigate the lamina-dependent disruption and recovery of the 3D architecture of the *Drosophila melanogaster* genome at topologically associating domains (TADs) resolution (~ 100 kb), using a model of the entire nucleus; weakening of chromatin–lamina interactions mimics a loss of chromatin organization associated with aging. We characterize this loss using the Shannon entropy of the appropriately normalized Hi-C contact and by deterioration of the chromosome territories.

Our main finding is that increase in Hi-C map disorder and deterioration of chromosome territories induced by lamina depletion are largely reversible when the wildtype (WT)-like Lamina-associated domain (LAD)–nuclear-envelope interactions are restored; the original and recovered conformational states of chromatin are essentially indistinguishable by the bulk Hi-C contact matrix, and the recovered WT state of chromatin remains stable. The recovery is robust: we find that the WT-like 3D architecture can also be recovered from a more severely perturbed lamina-depleted and compartment-disrupted state, in which all attractive interactions have been switched off.

The direct experimentally testable prediction is that the restoration of functional LAD–lamina interactions will promote recovery of the WT-like 3D chromatin architecture after lamina-dependent architectural disruption. We further hypothesize that lamin restoration may increase lifespan of the fruit fly and improve other aging-associated phenotypes through the recovery of young-like LAD–lamina attachments.

## 1 Introduction

Aging is a remarkably complex, multifactorial phenomenon that remains one of the grand unsolved problems of biology and medicine. Despite decades of research, there is no universally accepted theory of why organisms age or how this process is regulated [70, 13, 35]. Numerous mechanisms have been proposed, ranging from the accumulation of molecular damage to dysregulation of signaling pathways and loss of tissue homeostasis [53, 38]. Theories of aging can be divided into two broad, non-mutually exclusive groups: error-based and program-based theories[4]. Error-based theories suggest that aging results from the accumulation of molecular and cellular errors and damage that repair mechanisms cannot fully counteract. In contrast, program-based theories suggest that aging is regulated by developmental genetic programs that, over time, become detrimental [24, 33], implying that the aging process emerges from genetic programs shaped by natural selection. Program-based theories of aging include the neuroendocrine theory [5] and the software design flaw hypothesis [23]. Among the most widely discussed explanations is the view that aging results primarily from the gradual accumulation of DNA damage and mutations, which over time compromise genomic integrity and cellular functions [67, 40]. Defects in DNA repair are also known to accelerate aging, and various forms of genomic instability are hallmarks of aged cells [101]. However, accumulating evidence, including healthy aging of cloned sheep [107] and normal lifespan of mice cloned up to 20 times [113], suggests that DNA damage alone may not fully explain the reversible and coordinated aspects of aging observed at the cellular and organismal levels. Recent work[106] points to a complementary or even alternative perspective that aging, at least on certain timescales, may arise from the progressive loss of epigenetic information rather than from the mere accumulation of genetic lesions[49]. In this recent “information theory of aging”, the primary driver of functional decline is viewed as the gradual erosion of the chromatin-based regulatory architecture that maintains cellular identity. Conceptually, the information theory of aging is close to error-based theories that focus on information degradation, while incorporating features of program-based theories, such as interaction with developmental regulatory pathways. Epigenetic characteristics — such as histone modifications, DNA methylation patterns, and higher-order chromatin organization in 3D — encode regulatory information beyond the genome sequence itself. Their dysregulation can lead to aberrant gene expression programs reminiscent of aged or senescence-like phenotypes [28, 36, 103] even if the underlying DNA sequence remains largely intact.

During aging, cellular identity often becomes “fuzzy”, with aged genomes being in some ways less stable and more disordered than young ones [47, 68]. The tendency of chromatin to move toward more disordered, higher-entropy states with the progression of aging is perhaps not surprising. Much more surprising is the striking partial rejuvenation of aged cells through transient expression of Yamanaka factors [83, 71], which supports the notion that at least some features of aging are epigenetically encoded in chromatin marks and potentially reversible. These results are consistent with the possibility that changes in chromatin organization can contribute causally to aging- or senescence-linked transcriptional programs, rather than merely reflecting them. The mechanistic details of how this kind of reversal is achieved remain unclear. The expression of reprogramming factors in somatic cells initiates targeted chromatin remodeling prior to transcriptional changes via genome-wide changes in the promoter H3K4 methylation and gains of active H3K4me2 enhancer signatures [54]. Ectopic expression of Yamanaka factors Oct4, Sox2, Klf4 *in vivo* restores youthful gene-expression patterns and epigenetic features in aged tissues, partially reversing functional decline [118]. The observed blurring of the originally sharp histone modification patterns demonstrates that the aging process corrupts epigenetic information, thereby increasing its Shannon entropy [118]. While in principle there is no contradiction with the second law of thermodynamics, as the cell is an open system, orchestrating a precisely fine-tuned reduction of disorder in a complex system is a non-trivial task (going the other way is, usually, much easier).

The range of questions that must be addressed before organismal rejuvenation is understood is immense, reflecting the complexity of aging itself. One key layer of epigenetic information that has not been widely explored in relation to the potential for age reversal, is the 3D organization of the chromatin. How can a disorganized, presumably high-entropy state of aged chromatin be restored to its functional, lower-entropy configuration? In physics, the restoration of a lower entropy state can be effected easily by an appropriately chosen external factor: *e*.*g*., freezing water into ice decreases the system’s entropy. Likewise, a magnetic field can act as an ordering agent, reducing randomness in the spin system and, thus, lowering its entropy as it drives the system toward a more ordered magnetic state. Not surprisingly, the situation in biology is expected to be much more complex[22, 117]. The folding–unfolding transition in proteins is a good example to illustrate the point [30]. Any protein can be easily brought to a high entropy, disorganized (denatured) state by placing it into a denaturing environment [30], such as high temperature, urea or acidic pH [85], yet only for small globular proteins of relatively simple topology [92] this transition is truly reversible, that is the protein can be brought back to its original low entropy functional (native) state by simply removing the denaturing environment. This simple manipulation often fails to restore the original 3D shape of larger or more complex proteins, which generally require additional mechanisms such as chaperones to fold properly. The functional 3D configuration/topology of chromatin is no doubt exceedingly more complex than that of any protein, so finding a way to reverse its age-related chromatin disorganization is expected to be highly non-trivial. Assuming that a reductionist approach may help here, as it did in the past with many other complex scientific problems, it is worthwhile to investigate simpler systems that have at least some of the key features of aging. To this end we note that the 3D chromatin organization is shaped by at least two major interaction types: (1) chromosome-chromosome interactions and (2) interactions between chromosomes and nuclear lamina, a dense meshwork of Lamin proteins underlying the inner nuclear membrane. Chromosome-chromosome (or chromatin) interactions are studied using Hi-C, a technique based on Chromosome Conformation Capture (3C) technology [63]. As a step toward a minimal model of aging-associated 3D genome disorganization, we focus here on interphase chromatin and on lamina-dependent architectural changes. Recent Hi-C experiments [108] demonstrated that such “interphase aging” of post-mitotic cells is meaningful: the experiments have revealed, in great detail, how chromatin organization in neurons evolves with age, clearly demonstrating significant changes in the chromatin structure in aging nuclei. When the brains of newborns, one-year-olds, ten-year-olds, thirty-year-olds, and eighty-year-olds human individuals were compared, ultra-long-range chromatin contacts and specific inter-chromosomal contacts were identified as markers of cerebellar granule cell age. It has been hypothesized that the restructuring of the 3D genome that occurs over the course of a human lifetime may be stabilized by cell-type-specific gene transcription, which eliminates the expression of unnecessary genes and transposable elements. Note that cell division leads to only partial rejuvenation, because each division introduces inevitable DNA replication errors, mitotic errors, and other forms of molecular damage[59, 69]. Moreover, the Hayflick limit imposes an additional constraint on how far this natural rejuvenation can extend in a normal (non-cancerous) somatic cell [37]. We argue that a full mechanistic understanding of aging and rejuvenation processes during interphase — including epigenetic remodeling of chromatin interactions [108, 55] — could inform tissue rejuvenation strategies that avoid errors associated with repeated cell division. An important caveat here is that it is still unknown whether architectural rescue alone is sufficient for tissue rejuvenation.

Another prominent manifestation of cellular aging is the loss of integrity of the nuclear lamina and the detachment of chromatin domains from the nuclear periphery. The nuclear lamina provides both mechanical stability for the nucleus and an organizing scaffold for chromatin. Lamina-associated domains (LADs) represent large heterochromatic genomic regions that interact with the lamina and contribute to spatial segregation of heterochromatin and euchromatin. The interactions of LADs with the lamina are identified by the DamID method and their nuclear position can be traced by microscopy [111, 110]. With aging – and more dramatically in premature aging disorders such as Hutchinson–Gilford progeria syndrome (HGPS) -— the structural coupling between the chromatin and lamina weakens. The resulting reorganization of chromatin is correlated with large-scale transcriptional dysregulation and changes in histone methylation patterns [76, 105]. In HGPS, mutations in the Lamin A (*LMNA*) gene lead to the production of progerin, a defective Lamin A isoform that compromises lamina integrity, linking nuclear envelope dysfunction directly to accelerated epigenetic aging.

Fruit flies (*Drosophila melanogaster*) have a short lifespan, a small genome, and extensive genetic tools, making them a useful and simpler model for studying aging [34]. Unlike mammals, which have three lamin genes (*LMNA*, LMNB1, LMNB2) and multiple splice variants, *Drosophila* has only two lamin genes (*LamC –* A-type lamin gene and *Dm0 –* B-type lamin gene), which makes fruit flies ideal for studying the connection between lamin proteins and aging. Multiple studies have demonstrated that both A-type and B-type Lamins in *Drosophila* can vary with age depending on tissue, and that such variation correlates with decline in tissue function, chromatin destabilization, or genomic instability[52, 61, 114, 15, 58, 31]. Experimental knock-down of *Dm0* in neurons led to a shorter lifespan, progressive motor impairment, and a loss of dopaminergic neurons in the protocerebral anterior medial cluster of the *D. melanogaster* brain [86]. The Lamin B mutation causes heterochromatin decompaction and transposon activation in the Drosophila germline, which is similar to what occurs during physiological aging [80]. This indicates a close link between heterochromatin maintenance at the nuclear periphery and aging mechanisms. The ectopic expression of Lamin B in dopaminergic neurons in the anterior medial region of the protocerebrum improves male flies’ locomotor activity but with little impact on their stress responses or lifespan [64]. Together, these studies indicate that Lamin depletion is a reasonable model of age-related changes in chromatin structure. The causal role of lamina–chromatin interactions in maintaining nuclear organization is well known [12], and has also been probed computationally[27]. In particular, recent models based on polymer physics [16, 109, 25] have demonstrated that disruption of lamina attachment sites can recapitulate experimentally observed chromatin remodeling that follows Lamin depletion or degradation [110, 10]. More directly in the context of aging, Brownian-dynamics simulations of a single human chromosome demonstrated that altered heterochromatin–lamina and heterochromatin–heterochromatin interactions can drive senescence- and progeria-like chromosome reorganization [16]. These models underscore the strong physical coupling between LADs and the nuclear envelope, suggesting that the 3D architecture of chromatin is not only biologically regulated but also mechanically stabilized by its interactions with the lamina[51].

A central unresolved question in epigenetic rejuvenation is whether the aged 3D architecture of the genome can, in practice, be returned to a youthful architectural state. If epigenetic age reversal is to restore normal cellular function, it is difficult to imagine a truly rejuvenated nucleus retaining an aged, disorganized 3D genome architecture. However, recovery of a complex spatial organization is not guaranteed. Even the refolding of medium-sized proteins can be slow, incomplete, or dependent on chaperones, despite the fact that the native fold is encoded by a single polymer sequence. The interphase genome is vastly larger, confined, topologically constrained, and organized only statistically rather than as a single fixed structure. There is an additional reason why such recovery is not guaranteed. The functional interphase architecture of large eukaryotic chromosomes is not generally expected to be an equilibrated polymer state. For example, chromosome territories were argued to likely arise as a kinetic consequence of decondensation from compact mitotic chromosomes, and estimated global equilibration times of about 5 years for *Drosophila* chromosomes and about 500 years for human chromosomes, far exceeding the relevant interphase timescales [98]. Subsequent polymer-physics and data-constrained 3D genome models have emphasized the same general picture: interphase chromosome organization reflects a combination of slow topological relaxation, chromosome confinement, sequence-dependent interactions, and chromosome–nuclear-envelope contacts, rather than rapid equilibration to a unique mixed state [46, 89, 29, 6]. Therefore, restoring the WT interaction landscape does not necessarily restore the WT interphase conformational ensemble: the biologically relevant state may be metastable[16], history-dependent[16], and constrained by the path taken out of mitosis.

Thus, an important physical question – and the main focus of this work – is whether age-associated disruption of chromatin architecture produces kinetically inaccessible states, or whether a young-like, far-from-equilibrium architectural ensemble can be recovered on biologically meaningful timescales when the appropriate organizing interactions are restored. Here we test this idea in a coarse-grained, experimentally grounded model of the *Drosophila* interphase nucleus. In this model, age-associated architectural degradation is represented by weakening chromatin–lamina interactions, a perturbation motivated by the known relationship between lamina dysfunction, heterochromatin disorganization, and aging/progeroid phenotypes. We then ask whether restoring the youthful interaction landscape is *sufficient* to drive spontaneous recovery of the young-like chromatin ensemble. An important technical step toward answering this question is the availability of a useful metric for aging-associated changes in 3D chromatin organization. Although epigenetic aging clocks based on DNA methylation patterns are widely used as age estimators[41, 83, 71, 77], they do not capture the full complexity of cellular aging. Ultimately, more comprehensive age prediction models should include the degree of deterioration of the 3D architecture of chromatin inside the cell nucleus. Recent studies suggest that at the higher organizational level, the nuclear structure and spatial organization of chromatin can be explored as new estimators of aging. For example, the morphology of cell nuclei has been studied as a biomarker of aging [88] and has been used to model cellular age with the help of an interpretable deep learning approach [90]. Other studies demonstrated that the 3D genome organization changes dramatically during the cellular aging process [108, 55], further supporting the idea that aging clocks that incorporate key aspects of chromatin architecture can be developed. Here, we take a step in this direction by rigorously defining the entropy of HiC matrix and exploring its properties in the relevant context.

## 2 Methods

### 2.1 Entropy of Hi-C matrix

The concept of entropy has been used in the field of chromatin data analysis and simulation [88, 119, 94, 56]. Specifically, in the context of epigenetics and aging, Shannon entropy was used to quantify genome-wide methylation stochasticity; a link between increased entropy and epigenetic aging was suggested[45]. Recently, stratified DamID signals were used[66] to define a relevant entropy quantity, which was correlated with aging: the orderly distinction between genome regions that normally associate strongly versus weakly with the nuclear periphery becomes blurred during aging and in the progeroid models.

Here we seek the simplest possible, yet physically meaningful, entropy-based metric that can be computed from a Hi-C contact matrix, while retaining all locus-locus interaction information at the chosen resolution. Specifically, we introduce the notion of entropy *S*(*A*_*ij*_) of a contact (Hi-C) matrix {*A*_*ij*_} that describes contact frequencies between *N* structural elements (loci) *i* = 1, 2, … *N* of a chromatin (polymer) chain. Naturally, we assume *A*_*ij*_ ≥ 0. Next, *A*_*ij*_ needs to be converted into a probability distribution *P*_*ij*_, to which the usual Shannon form, *S* = − ∑_*ij*_ *P*_*ij*_ log *P*_*ij*_, can be applied, with all expected properties of *S*, such as that it is maximized by a uniform underlying probability distribution. There is more than one way to convert *A*_*ij*_ into *P*_*ij*_. Since our goal is to derive an expression for *S* that treats all “non-trivial” elements of the contact matrix on the same footing, as opposed to, *e*.*g*., focusing on near-diagonal elements [56], there are three intuitive ways to normalize *A*_*ij*_: global normalization 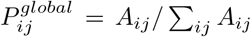, as well as row-based and column-based normalizations 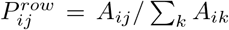 and 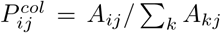, respectively. Once the normalization is chosen, we can define

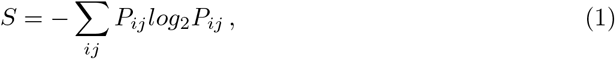

where we have chosen *log*_2_ so the entropy is expressed in bits. This quantity is the Shannon entropy[104] of the row-normalized contact-probability distribution, and should not be confused with the thermodynamic or conformational entropy of a polymer defined by counting conformational states consistent with macroscopic constraints[87, 99].

The following limiting, albeit hypothetical, cases help choose the best normalization strategy and justify the definition, Eq. 1, specifically in the context of 3D organization of chromatin. On the one hand, a completely disorganized contact map would have the same *P*_*ij*_ values for all its elements, which means that every locus has an equal probability of being in contact with every other. Presumably, this would be a *maximum-disorder* state [9] of chromatin of maximum entropy. On the other extreme is a purely diagonal matrix of contacts: *P*_*ii*_ = 1, and *P*_*ij*_ = 0 for all other elements outside the diagonal, *i* ≠ *j*. Physically, this contact matrix would describe chromatin without any interactions except between the neighboring loci along the chain. All real biological cases are in-between: the off-diagonal contacts *P*_*ij*_ *>* 0. While the global normalization – a single normalizing constant for all the elements – has the appeal of being most intuitive, as well as of preserving the expected symmetry *A*_*ij*_ = *A*_*ji*_ of the contact matrix, the global normalization has a fundamental drawback, as it violates entropy additivity for non-interacting systems. Namely, the entropy defined via globally normalized matrix is non-additive for independent, non-interacting subsystems, such as two independent, non-interacting chromosomes. Such systems are described by block-diagonal matrices, and one easily sees that the global normalization of the elements in each block depends on the presence and size of the other block(s), even if all the “cross” elements between the blocks are strictly zero. Fortunately, either the row-based or column-based normalization is free from this critical defect: the entropy of a block-diagonal matrix is strictly the sum of entropies of its blocks. Moreover, while the resulting probability matrices *P*_*ij*_ are no longer symmetric, the symmetry is effectively restored as one takes the sum in Eq. 1: *S*^*row*^ = *S*^*col*^. This is because, for a symmetric *A*_*ij*_, 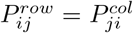, that is, the probability vector that defines a row in the row-normalized matrix is exactly the probability vector that defines the corresponding column in the column-normalized matrix, just transposed. We therefore choose the row-based normalization scheme for the rest of this work: *S* = *S*^*row*^. As defined, 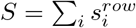, where 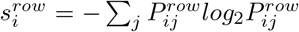; for the rest of this work we drop the label “row” for notational simplicity.

Note that with this definition, *S* grows proportionally to the total number of elements *N*; where relevant, and to facilitate comparison between contact matrices of different sizes, one can also consider the average entropy per row, *S/N*. Since our primary use of the above defined Hi-C entropy here is to trace temporal evolution of chromatin disorder for each system separately, in what follows we do not explicitly report *S/N*. In what follows, we will be excluding *K* near-diagonal elements, |*i* − *j*| ≤ *K*; for small *K* ≥ 1 the procedure yields an entropy measure that is sensitive to medium- and long-range structural features of chromatin, while removing trivial polymer connectivity effects and de-emphasizing very short range structural elements. Note that for a connected chain, *A*_*ii*_ = 1 and *A*_*i,i*+1_ = 1 always, so it makes sense to exclude these elements from the calculations to remove this trivial bias towards the diagonal; in practice, these elements are set to zero (masked) prior to the normalization procedure. For simplicity and ease of interpretation, we use *K* = 1 as the default, which removes self-contacts and immediate polymer-neighbor contacts while retaining near-local chromatin organization beyond trivial chain connectivity. The effect of larger near-diagonal masks (*K >* 1) is examined in the SI. The central conclusions that in both the *lamina-depleted* and *lamina-depleted + compartment-disrupted* states, contact-map entropy is increased relative to WT, and that restoration of the WT interaction landscape returns the entropy toward WT values, are unchanged when *K* = 4 is used. The more subtle relative ordering of the *lamina-depleted* and *lamina-depleted + compartment-disrupted* states is *K*-dependent, and is discussed in the SI.

Also note that, as expected, *S*(*A*_*ij*_) depends on the level of coarse-graining (resolution) of matrix *A*_*ij*_ – direct comparison of entropy values obtained for different genomes or the same genome at different resolution should be avoided. Unless otherwise specified, all Hi-C matrices used in the work are coarse-grained to a resolution of 100kb. A more subtle but related feature of *S*(*A*_*ij*_) is that *S*(*A*_*ij*_) depends on the averaging performed on *A*_*ij*_: averages over more states or longer times, in the case of time-dependent *A*_*ij*_, are expected to result in more “spread out” matrices, and hence higher *S*(*A*_*ij*_). Therefore, most meaningful comparisons should be limited to *A*_*ij*_ averaged over the same time intervals and the same number of states/topologies sampled.

For chromatin structures made up of several distinct and unconnected chromosomes, because Hi-C matrices are block-diagonal at the chromosome level, we exclude (mask) shortrange contacts |*i* − *j*| ≤ *K* only *within* each chromosome block, so that the exclusion does not affect contacts between different chromosomes. All practical details of extraction (in the case of Progeria Hi-C maps), coarse-graining, and post-processing of Hi-C matrices, including normalization, exclusion of near-diagonal elements, entropy evaluation, and visualization are described in the corresponding sections of the SI.

### 2.2 Processing of experimental Hi-C maps

#### The WT vs. Progeria maps

We started from the original deposited[76] Hi-C maps of (Father-p18) and progeria (HGPS-p19) chromosome 7 matrices from GSE41763. To reduce sensitivity to the substantial sparsity of the experimental Hi-C maps, we have coarse-grained the matrices. At the original 200-kb resolution, 61.2 % and 86.4 % of the finite matrix elements were zero, respectively. After trimming both maps to a common genomic interval, each complete non-overlapping (5 × 5) block of 200 kb matrix elements was summed, yielding matched (158 × 158) matrices at 1 Mb resolution. This procedure reduced the zero fractions to 1.4 % and 8.8 %, respectively, which we have deemed acceptable. We then applied the |*I* − *j*| ≤ *K* = 1 exclusion and row-normalization as described above. For visualization, the row-normalized contact probabilities were displayed on a logarithmic color scale.

#### The WT vs. lamina depleted maps

We started from the original deposited[110] Hi-C maps of WT and lamin knockdown fruit fly nuclei (whole genome). For consistency with the topologically associating domain (TAD)-resolution computational model of the fruit fly nucleus used in this work, the matrices were coarse-grained to the specific partitioning of the genome into TADs employed here is described in Refs.[60, 109]. Each complete non-overlapping block of the original Hi-C map that fit within the boundaries of each TAD was summed; summation over the two experimental replicates was also performed. We then applied the |*i* − *j*| ≤ *K* = 1 exclusion and row-normalization as described above. For visualization, the row-normalized contact probabilities were displayed on a logarithmic color scale.

### 2.3 Chromosome Intermingling

Inter-chromosomal intermingling is quantified as the fraction of the chromatin-occupied nuclear volume that is shared by two or more chromosome territories, by analogy with the cryo-FISH “intermingling volume” described in Ref. [11]. Here, each of the six major chromosome arms of the fruit fly (2L, 2R, 3L, 3R, 4 and X) is treated as a distinct territory.

For every conformational snapshot, the bounding box is discretized onto a regular cubic grid of spacing d*x* = 0.1 *σ* (*σ* = simulation length unit, which is 1 *µ*m in this work). Each monomer is assigned a spherical footprint of radius *R* = 0.2 *σ*: a voxel is marked as occupied by an arm if its center lies within *R* of any bead of that arm, and the union of such voxels defines the arm’s territory. A voxel is classified as *intermingled* when occupied by at least two of the six arms. The per snapshot (per frame) intermingling index is computed as

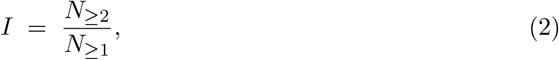

where *N*_*≥*2_ and *N*_*≥*1_ are the numbers of voxels occupied by at least two and at least one arm, respectively; *i*.*e*. the fraction of the nuclear volume occupied by the chromatin in which territories overlap. By definition, *I* is dimensionless, bounded in [0, 1], and independent of the grid origin and of the *σ*→ *µ*m calibration. The intermingling index of fruit fly chromatin is shown in Figs. 5 and 7. Further details and a connection with Ref. [11] are available in the SI.

### 2.4 The dynamic model of the fruit fly nucleus

Our approach uses the well-established computational strategy [98, 60, 109, 65, 84, 57, 93, 109, 78, 100, 20, 48, 62, 17, 29, 26, 19] to represent chromatin at an appropriate coarse-grained level. Specifically, we employ a representation of the fruit fly genome at the level of TADs[109], *i*.*e*. ~ 100 kb of chromosome length. Each TAD is modeled as an interacting polymeric unit whose effective attractions reflect the underlying chromatin state and affinity to the lamina. This level of abstraction captures both the essential large-scale organization of interphase chromatin and the stochastic thermal dynamics that underlie its remodeling. The model allows systematic variation of physical parameters—such as the strength of lamina–TAD interactions, inter-TAD contact potentials, and nuclear confinement—to reveal the energetic and entropic principles governing chromatin reorganization. The choice of TAD level resolution is well justified for the main purposes of this work: a recent experimental study has demonstrated that lamina depletion affects only high-level chromatin organization, including chromosome territories, chromatin compartments and giant chromatin mega-loops while the TAD structure and loops inside TADs remain intact[116].

Except for the experimental Hi-C maps shown in Fig. 3, the 3D structures of fruit fly chromatin and their time evolution described in this work were generated using the model fully detailed in Ref. [109] and recently employed in, *e*.*g*., Refs. [1, 74]. Briefly, the model describes the time evolution of embryonic *Drosophila* female interphase nuclei at spatial resolution of topologically associating domains (TADs). In the model, a diploid set of four female chromosomes is represented by four chains of spherical beads. Each bead accounts for a pair of homologous TADs in paired homologous female chromosomes; the beads are of variable size, the average radius of the bead is 0.091 *µ*m. In addition to 1169 TAD-beads, the chromosome chains also contain the pericentromeric constitutive heterochromatin (HET) and centromeric (CEN) regions, represented by additional 6 and 4 beads, respectively, and an almost centrally located nucleolus. The chromosome chains and the nucleolus are confined to a spherical enclosure representing the nuclear envelope (NE), the inner surface of the nuclear envelope is lined by the nuclear lamina (NL) – a meshwork of Lamins. A supplemental movie is available that illustrates ~ 30 seconds of the time evolution of the model nucleus.

The model agrees with experimental data on multiple independent metrics [109]: the Pearson’s correlation coefficient of the model-derived Hi-C map with the experimental wild type (WT) Hi-C map is 0.954; reasonably distinct chromosome territories 11 hrs into the interphase; model-derived radial chromatin density distribution is in good agreement with the experimental one. The model is also consistent with the experimentally observed stochasticity and cell-to-cell variability of the attachment of LADs to the NE.

The interaction parameters of the models were tuned to reproduce the TAD-NE contact probability data [91] and the coarse-grained TAD-TAD contact probability matrix, derived by Li et al. [60] from the experimental bulk Hi-C map [102]. The predominant types of TAD-TAD interactions are active TAD–active TAD and inactive TAD–inactive TAD. These attraction result in the segregation between the A- and B-compartments. Each TAD-related bead in the models represents two paired homologous TADs with approximately 100 kb of DNA in each, resulting in one polymer chain for each of two paired homologous chromosomes in the diploid female *Drosophila* genome. These homologous TADs and chromosome pairing are observed experimentally [32, 21, 2, 18], and explained within a theoretical model [18]. The model fully accounts for the different experimentally observed topologies (mutual arrangements of the chromosomes [39]) of the *Drosophila* chromatin. Unless otherwise specified, all of the model-derived quantities, including the bulk Hi-C maps *A*_*ij*_, the entropy *S*(*A*_*ij*_), and the chromosome intermingling are calculated as time averages, averaged over all 18 trajectories generated by the model. The relation between the experimental and simulation timescales was previously established by comparing the calculated and experimental mean square displacement (MSD) of a chromosomal locus as a function of time; see Ref. [109] for details.

Four initial configurations of the chromosomes – mutual arrangement (topology) of the chromosome chains – are considered[109]; to account for the different experimental weights of these arrangements, we used two identical replicas for two of them, making the total number of properly weighted chromosomal arrangements equal six. To account for the natural variability of nucleus size in different nuclei, 3 slightly different nucleus sizes (2 ± 0.05*µm*) are considered for each topology [109]. Thus, as in our previous work, the entire simulated ensemble consists of 18 model nuclei. Langevin dynamics is used to simulate the resulting 18 distinct model nuclei and generate 18 trajectories, each representing hours of the biological time evolution of the interphase chromatin, *e*.*g*., Figs. 4 and 6. In this work, each nucleus is simulated to produce three time intervals of about 3 hrs. of biological time each.

Each data point in Figs. 4 and 6 is an average over 10 minutes, or 6065 conformational snapshots, except for the inset Hi-C maps, which are averages over 3 hrs, or 109182 snapshots. The average intermingling, ⟨*I* ⟩, is computed over 1000 equidistant snapshots representing each state of chromatin (3 hrs of simulation), see SI.

The simulations reported in this work took approximately 20 days on 18 modern computing cores.

#### 2.4.1 *lamina-depleted* model nucleus

Lamin depletion is modeled by setting the affinity of LAD-containing TADs for the NE to zero, see Ref. [109] for details. In particular, it was shown previously [109] that the model reproduces well the changes in chromatin distribution induced by Lamin depletion. The selection of the representative ensemble is the same as for the wild type (WT) nuclei described above. We refer to this model state as *lamina-depleted*.

#### 2.4.2 *lamina-depleted + compartment-disrupted* model nucleus

In this hypothetical nucleus, all attractive interactions are switched off, including LAD-NE (as in the *lamina-depleted* state), all TAD-TAD, and the attraction to the nucleolus. The model is therefore reduced to self-avoiding chain(s) under confinement. The selection of the representative ensemble is the same as for the wild type (WT) nuclei described above.

### 2.5 Software used

VMD [43] was used to visualize chromosome models, *gnuplot* to make graphs, Juicebox[97] to visualize and pre-process Hi-C maps. ChatGPT versions 4 and 5 was used to generate and analyze FJC ensembles, explore various options for Hi-C entropy normalization in simplified cases, and help with grammar and style editing. The movies were generated with the help of Claude Code (Anthropic), using the Claude Opus 4.8 model. The authors assume full responsibility for the content of this manuscript.

## 3 Results

Using a realistic model of 3D chromatin organization in fruit fly, we test whether lamina-dependent architectural disruption that mimics age-associated chromatin disorganization can be reversed by restoring the interaction landscape characteristic of the WT nucleus, despite the complexity and highly dynamic nature of chromatin organization. To follow this process quantitatively, we use two complementary descriptors of 3D genome organization within the nucleus: Hi-C contact-map entropy introduced in this work and chromosome intermingling[11]. We begin by introducing Hi-C contact-map entropy and show that it traces the expected deterioration of 3D chromatin architecture, and is relevant to age-related changes in the 3D organization of chromosomes.

### 3.1 Shannon entropy of Hi-C contact map as a metric of chromatin disorder in the interphase

In what follows, we approximate the 3D chromatin organization by its corresponding contact probability matrix *P*_*ij*_, which, in turn, can be approximated from a suitably normalized Hi-C map. From the examples discussed in the introduction, one can generally conclude that, at least for a single chromosome, the contact matrix becomes more “disordered” with age, or with the onset of conditions, such as Progeria, that can be considered as models of aging. Specifically, one expects that the “older” contact maps will have more distant off-diagonal contacts [108] compared to the younger nuclei, at least in the interphase, which is the focus of this work. To capture this type of disorder in a Hi-C matrix *A*_*ij*_, we propose to use the familiar concept of entropy:

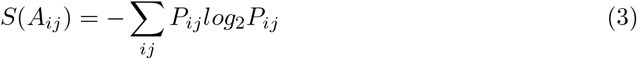

where we have chosen *log*_2_() to specify Shannon entropy [104], as opposed to *ln*() for Gibbs entropy [87] standard in Physics, for reasons mentioned in Methods. All important technical details, including the choice of normalization condition needed to convert the experimental Hi-C matrix to the matrix of contact probabilities *P*_*ij*_, are presented in Methods.

The behavior of the proposed definition of Hi-C entropy is illustrated on several simplified and limiting cases, going from completely hypothetical to more biologically realistic, Fig. 1. First, we consider a completely uniform distribution of contacts *A*_*ij*_ = *const*, Fig. 1(a), corresponding to the completely hypothetical case of a *maximum-disorder* state, in which every locus has an equal chance to come in contact with any other locus, regardless of the inter-loci distance. Next, we consider a more realistic case of two block-diagonal matrices with added non-zero elements between the blocks, corresponding to several chromatin chains that interact, Figs. 1(b,c). For relevance to this work, the number of blocks and the length of each block are set equal to the number of chromosomes and the number of TADs in each chromosome of *D. melanogaster*, respectively, summing to the same total number of TADs [102, 109], see “Methods”. For simplicity, we assume uniform distributions of contact probabilities within each block *P*_*B*_, and between blocks *P*_*I*_, and vary their ratio *P*_*i*_*/P*_*B*_. Finally, in Fig. 1 (d), we consider a model of a single chromosome, whose pairwise contact matrix is that of a single freely jointed chain (FJC) of the same length as the entire genome of fruit fly, at TAD resolution, see “Methods” for details. In all cases, the total number of matrix elements *N* × *N* = 1169^2^ is the same; the corresponding entropy values are summarized in Table 1, to which we have also added the trivial case of a completely diagonal contact matrix for which *S* = 0.

**Table 1:** Comparison of entropies of the four idealized Hi-C matrices described in Fig. 1. The last row is added for completeness.

| System | $N$ | $S$ (bits) |
| --- | --- | --- |
| (a) Uniform distribution | 1169 | 11909 |
| (b) Strong interactions | 1169 | 10894 |
| (c) Weak interactions | 1169 | 9540 |
| (d) Freely jointed chain | 1169 | 7746 |
| Fully diagonal contact matrix | 1169 | 0 |

**Figure 1:**
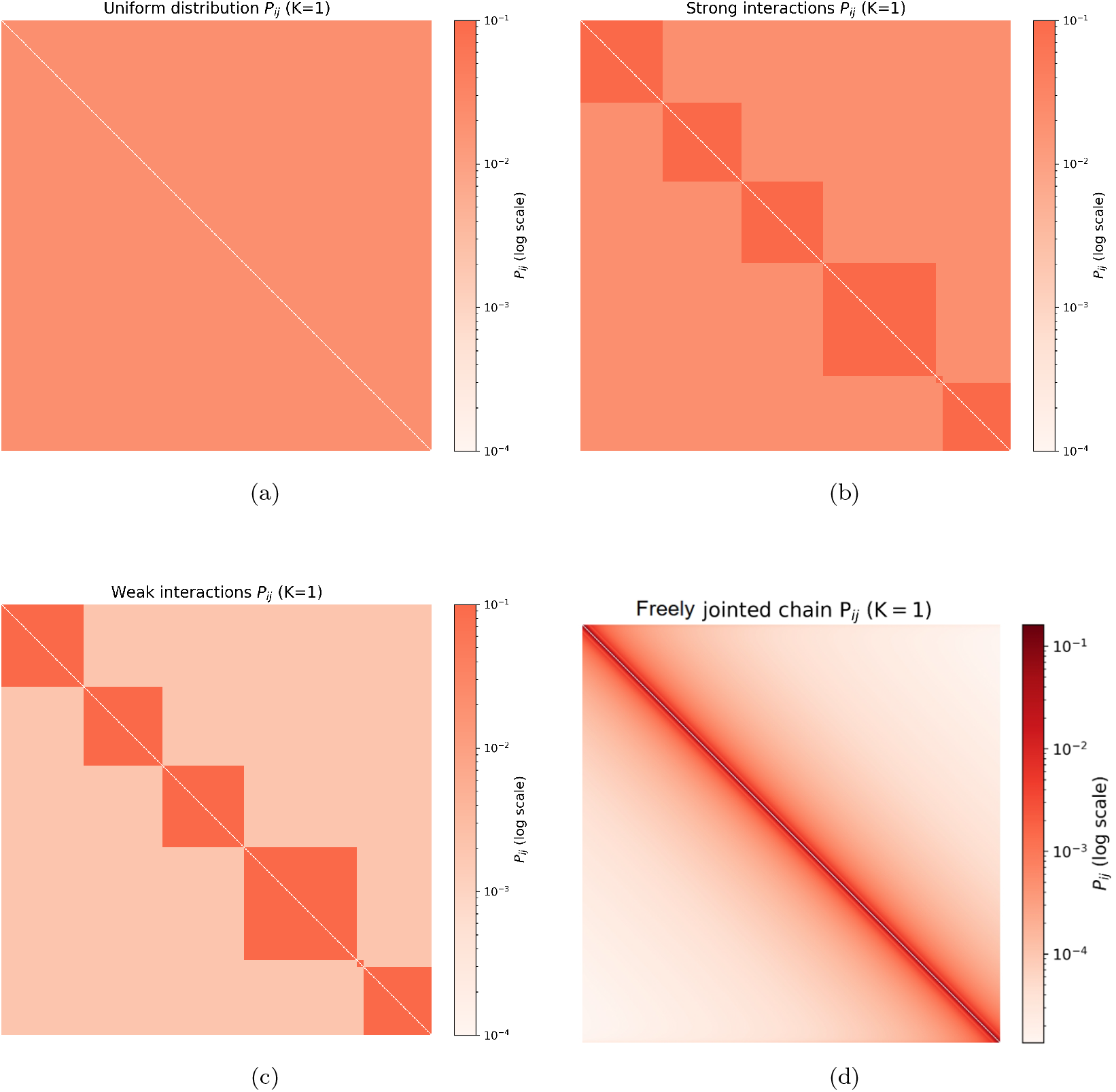
Examples of contact probability (Hi-C) square *N* × *N* symmetric matrices, *N* = 1169, corresponding to four idealized systems. (a) A *maximum-disorder* state – equal probability of all contacts; (b) Strong interaction between different “chromosomes” *P*_*I*_*/P*_*B*_ = 10^*−*1^; (c) Weak interaction between different “chromosomes” *P*_*I*_*/P*_*B*_ = 10^*−*2^; (d) A single freely jointed chain of *N* beads.

As expected, the proposed definition of Hi-C map entropy anti-correlates with the amount of “structure” in the Hi-C map, with the highest entropy corresponding, as expected, to a completely uniform distribution. In the SI a brief exploration of potential applicability of the proposed Hi-C entropy *S*(*A*_*ij*_) in the context of mammalian cell aging during interphase is presented.

#### 3.1.1 Hi-C entropy increases in premature aging (Progeria) associated with lamina dysfunction

As discussed in the Introduction, *lamina-depleted* is a reasonable model of age-related changes in chromatin structure. As an initial semi-quantitative consistency check, we compared WT and Progeria-associated Hi-C maps, Fig. 2. Hutchinson-Gilford Progeria syndrome (HGPS) is a premature aging disease that is often caused by a point mutation at position 1824 in LMNA. In-vivo, these changes can lead to loss of spatial chromatin compartmentalization and mis-regulation of transcription[76]. What is most relevant to us here is that this mutation disrupts heterochromatin-lamina interactions: similar *lamina-depleted* states can be realized in fruit fly, both experimentally[110, 10] and computationally[109, 1, 74, 51].

**Figure 2:**
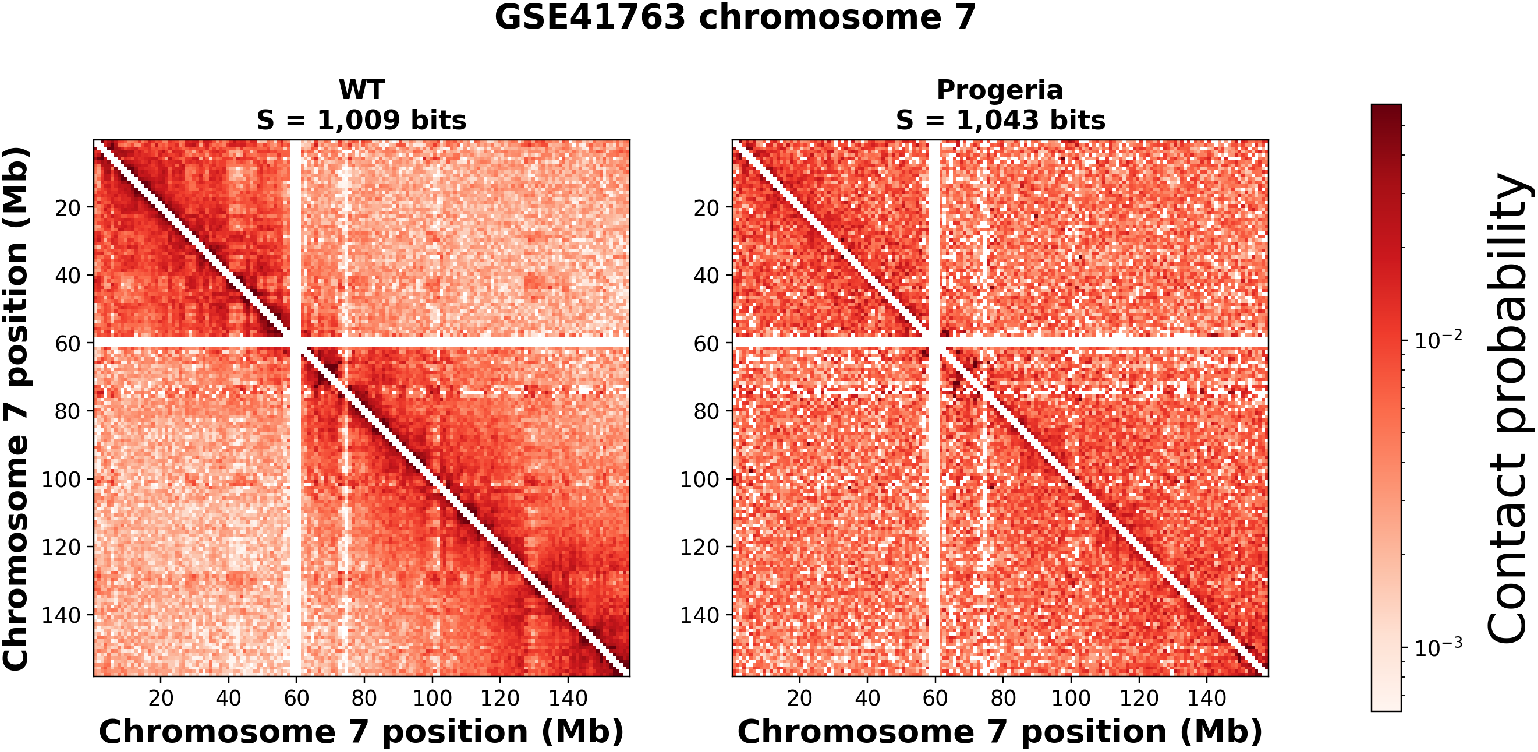
Row-normalized Hi-C contact probability maps for WT (Father-p18, left) and Progeria/HGPS (HGPS-p19, right) chromosome 7 fibroblast samples from Ref. [76]. The HGPS map is visibly more dispersed and less compartmentalized than the WT map and yields a higher Hi-C contact-map entropy, consistent with increased 3D genome disorder associated with lamina dysfunction in premature aging. The Hi-C entropy values are estimated from the deposited experimental contact maps as described in “Methods”.

As is evident from Fig. 2, the HGPS Hi-C map is visually more dispersed and less compartmentalized than the WT map, consistent with lamina-associated disruption of 3D genome organization. The Hi-C contact-map entropy is higher in HGPS than in WT, *S*(*WT*) − *S*(*Progeria*) = 34 bits. Thus, this comparison supports the expected direction of the entropy change in a progeroid context, whereas the Drosophila lamin-depletion data below provide a more directly relevant experimental comparison for the present model.

#### 3.1.2 Entropy of the Hi-C map increases with lamina depletion in fruit fly

Experimental and computational works show that lamina depletion at the nuclear envelope (NE) in fruit fly can lead to significant re-arrangements of chromatin, as evidenced by major shifts in the chromatin density distribution upon lamina depletion [10, 109]. However, the corresponding changes in the Hi-C map are much more subtle, see, e.g., Ref.[109], and Fig. 3. Therefore, it is reassuring that the proposed metric, *S*(*A*_*ij*_), Eq. 3, is able to reliably capture the subtle increase in the contact map disorder induced by experimental lamina depletion (lamin knockdown[110]). We stress that this specific data point applies to the whole interphase nucleus of fruit fly, which makes it most relevant for this work. This result supports the usefulness of Hi-C contact-map entropy as a metric for detecting lamina-dependent, aging-associated changes in chromatin organization at the scale of the fruit fly nucleus.

Another critical observation is that the values of Hi-C entropy of the *experimental* WT and *lamina-depleted* states, Fig. 3, are very close (within 1.6 %) to those calculated from Hi-C entropy maps of the WT and *lamina-depleted* states predicted by our computational model of fruit fly chromatin, see also the following section for the specific numbers. This close agreement with the corresponding experiment provides support for the computational model’s ability to capture the direction and magnitude of lamina-depletion-induced changes in Hi-C entropy.

**Figure 3:**
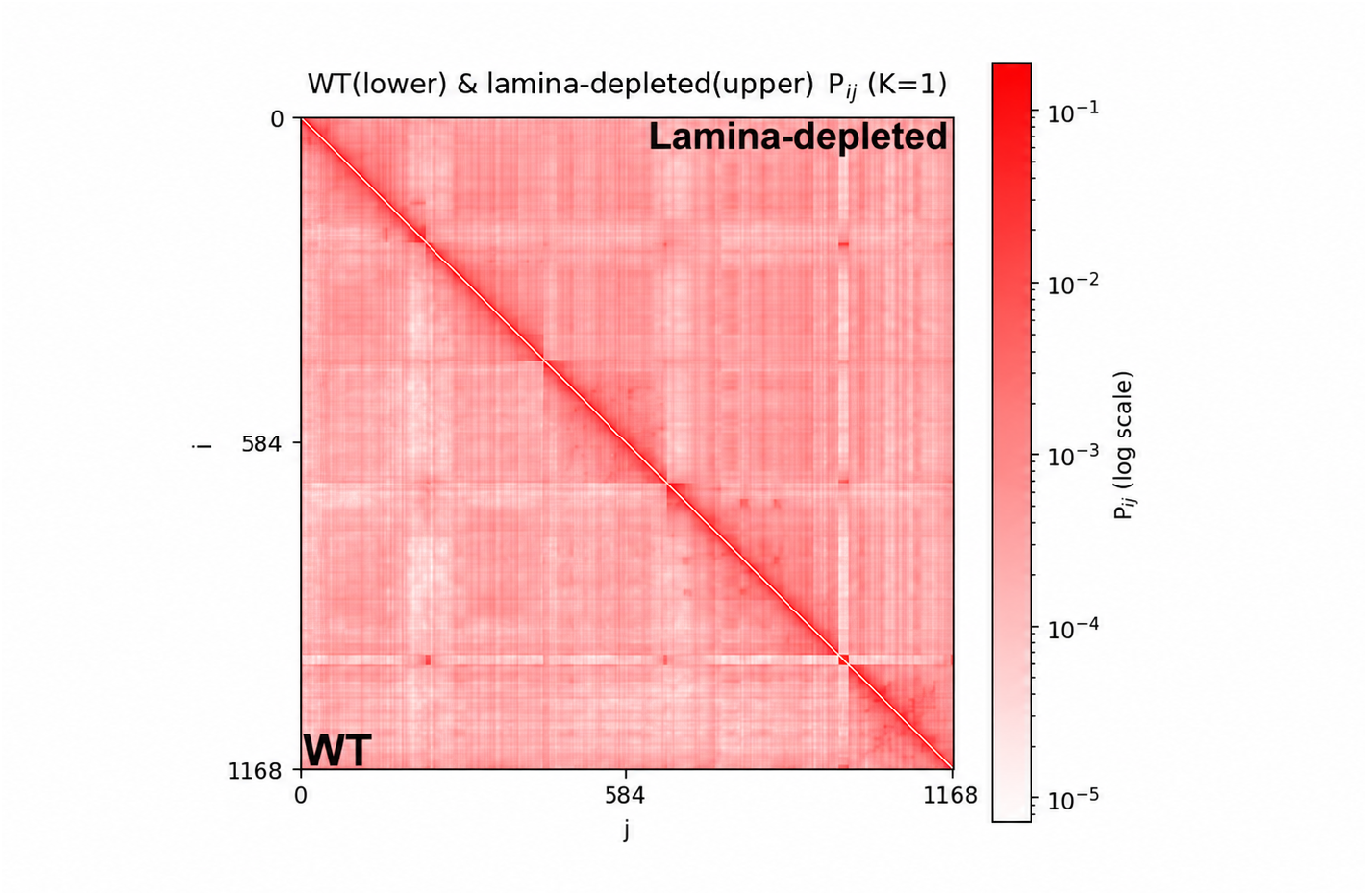
Changes in fruit fly chromatin architecture upon *lamina-depleted*. Shown are WT (lower triangle) vs. *lamina-depleted* (upper triangle) row-normalized contact probability map of the whole fruit fly nucleus at TAD resolution calculated directly from the experimental Hi-C data in Ref. [110]. The corresponding Hi-C entropy values are: *S*_WT_ = 8022 bits, *S*_LM_ = 8849 bits. The TAD index, which grows monotonically with the genomic coordinate, is shown along the axes. The average TAD size is about 100 kb; the specific partition of the fruit fly genome into TADs is described in Refs.[60, 109]

### 3.2 Restoration of a young/WT-like 3D state of chromatin in fruit fly

In this section, we address the key question of this work: can one reverse aging-associated disruption of the structural state of WT chromatin in fruit fly? In particular, to what extent can the youthful conformational ensemble of the chromatin be restored by eliminating/reversing the external factor or condition that has caused the deterioration of the original 3D structure of chromatin?

#### 3.2.1 Recovery from *lamina-depleted* state

The key result of this work is summarized in Fig. 4, which shows the temporal evolution of the chromatin organization of the fruit fly nucleus as it transitions from the WT to the *lamina-depleted* state, and then “rejuvenated” to the WT state. Specifically, after spending 3 hrs in its WT state, the chromatin undergoes an induced transition to *lamina-depleted* state, in which all the attractive interactions between LADs and the nuclear envelope have been switched off, see Methods. As expected, the transition causes the entropy of the corresponding contact matrix to increase substantially. Remarkably, the values of Hi-C entropy maps of the *computed* WT and *lamina-depleted* states, Fig. 4, are within 1.6 % of the *experimental* WT and *lamina-depleted* states, Fig. 3. The corresponding changes in Hi-C entropy upon lamina depletion, *S*(*lamina-depleted*) − *S*(*WT*), are also very close between the simulated and experimental Hi-C maps. This agreement with the relevant experiment further supports the use of our computational approach for generating experimentally testable predictions about lamina-dependent architectural recovery.

**Figure 4:**
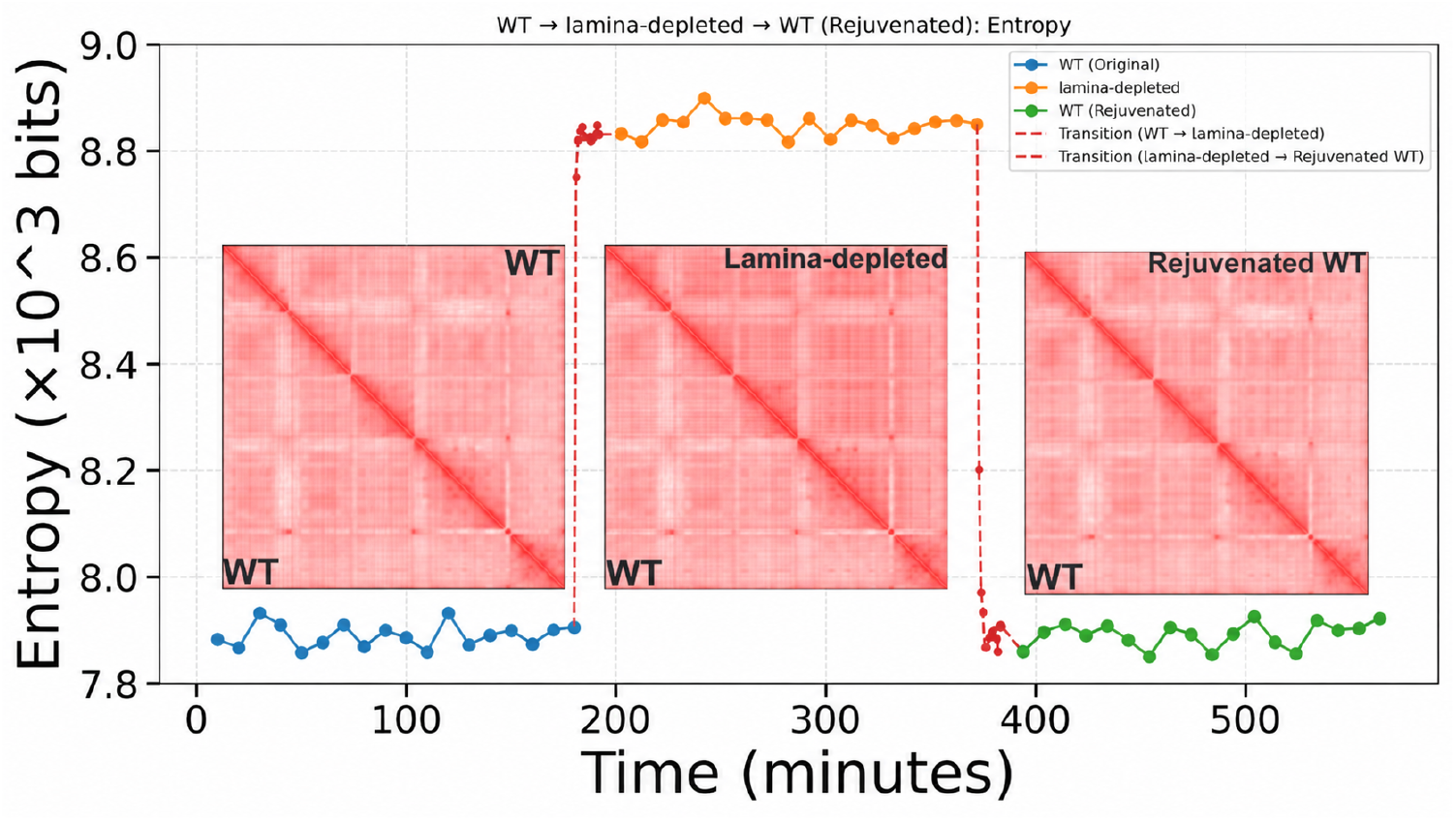
Transition from the WT to *lamina-depleted* state of fruit fly is followed by its “rejuvenation” that fully restores the attractive interactions between LADs and the nuclear envelope (NE), present in the WT state and absent from the *lamina-depleted* state. The transition to/from the *lamina-depleted* state is initiated by an instantaneous switching off/on of the LADNE attractions, see Methods. The corresponding Shannon entropy values, averaged over time, are ⟨*S*_WT_⟩ = 7889 ± 5 bits, ⟨*S*_Lamina depleted_⟩ = 8850 ± 6 bits, and ⟨*S*_Rejuvenated WT_⟩ = 7893 ± 6 bits, where the error bars represent the standard error of the mean. Each of the three Hi-C maps shown as the insets is a bulk Hi-C map averaged over the full 3-hour window corresponding to each of the three states: WT, *lamina-depleted*, and rejuvenated WT. To facilitate visual comparison, the lower triangle block of each matrix is the same original WT, while the upper triangle shows the state corresponding to the time interval.

The deterioration of the chromatin organization in the *lamina-depleted* state (upper triangle) is visible in the Hi-C map by comparing it with the WT map (lower triangle). That deterioration of the 3D organization of chromatin also manifests itself in substantial decay of chromosome territories[51], as evidenced by a two-fold increase in chromosome intermingling, Fig. 5. The decay of chromosome territories following chromatin detachment from the nuclear periphery due to lamina depletion is consistent with the original computational prediction made for both polytene and interphase chromosomes of fruit fly[50, 51], and has since been confirmed experimentally in human stem cells[116].

**Figure 5:**
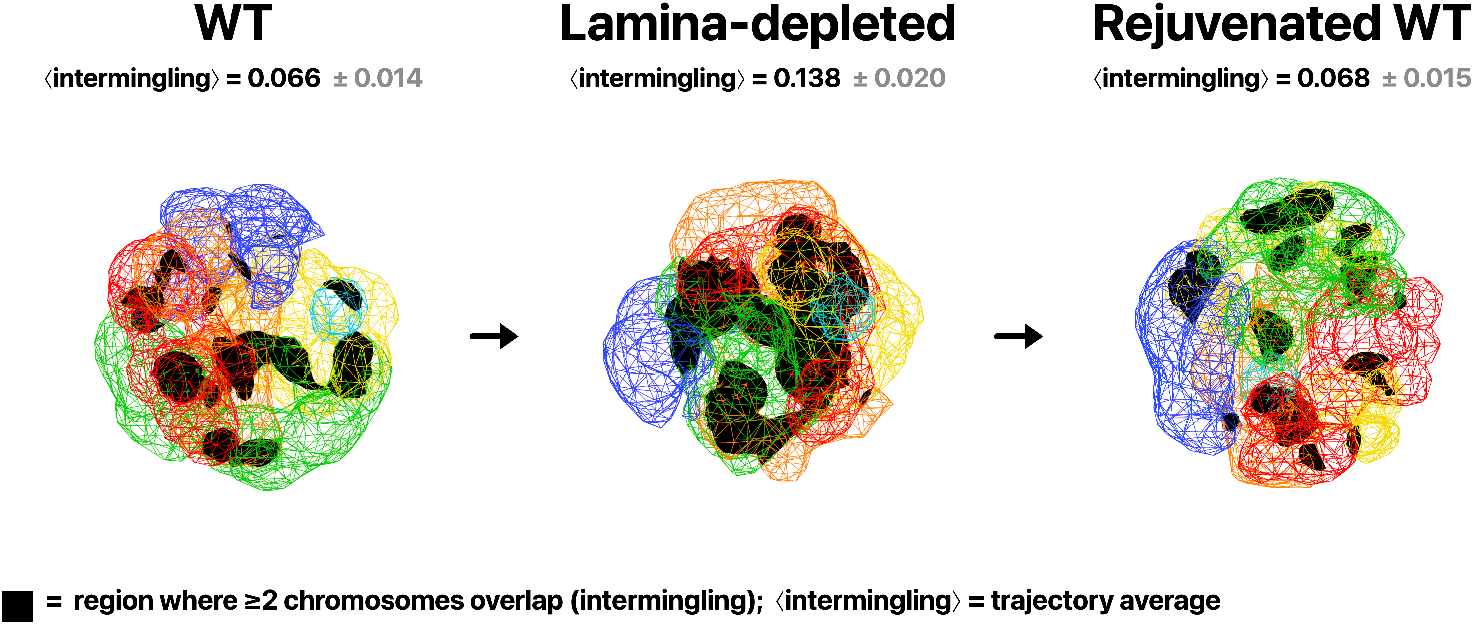
Chromosome territories along the WT → *lamina-depleted* → WT rejuvenation pathway. The intermingling between chromosomes increases following transition to the *lamina-depleted* state, and returns to its WT value upon the restoration of the attractive interactions between LADs and the nuclear envelope (NE), present in the WT state and absent from the *lamina-depleted* state. The snapshots shown are intended to illustrate each of the three states in Fig. 4; see also SI movies that show these states in dynamics. Each snapshot is selected so that its chromosome intermingling value closely approximates the trajectory average (shown above the snapshot) of the corresponding state. The intermingling regions are colored black. Each chromosome arm is colored differently, the gray sphere is the nucleolus. For this illustration, only one out of four experimentally observed mutual arrangements (topologies) of the chromosomes (CYSX-6S nucleus topology[109]) is shown. The errors are estimated as standard error of the mean.

After equilibrating for 3 hrs in the *lamina-depleted* state, the “rejuvenation” process is initiated: the attractions between LADs and the nuclear envelope, characteristic of the WT chromatin, are fully restored. Remarkably, the entropy of the resulting rejuvenated state quickly (within at most 10 minutes) returns to its WT value within statistical uncertainty, see the caption of Fig. 4. The recovered state is nearly indistinguishable from the original WT state by bulk Hi-C contact-map similarity, see the insets in Fig. 4. Qualitatively, the Hi-C maps of the two states are visually indistinguishable at the resolution of the model. Quantitatively, the contact probability difference map (not shown) between the two consists of matrix elements of the order of 10^*−*4^ or smaller, that is 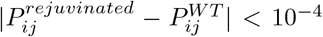. The Pearson correlation coefficient between the original WT and recovered maps is 0.999902. Consistent with nearly complete restoration of the WT-like Hi-C map in the rejuvenated state of chromatin, chromosome intermingling is reduced to its WT level, Fig. 5, reversing the partial decay of chromosome territories induced by the lamina depletion. We stress that the restoration of the WT state refers to the corresponding statistical ensemble, and not to its individual conformations, which are highly dynamic on the relevant time scales[109] (see also the supplemental movie).

Thus, to the extent that Lamin depletion mimics an age-associated deterioration of chromatin architecture in fruit fly, the model predicts that this deterioration can be reversed by restoring the LAD–nuclear-envelope interactions characteristic of the young WT nucleus^a^. Given the complexity and dynamic nature of interphase chromatin, this recovery is non-trivial: the model does not impose the positions of individual chromatin domains, but instead restores the physical interaction landscape that stabilizes the young architectural ensemble. The test is therefore not whether the WT state is thermodynamically favored after restoration of the WT parameters; rather, the test is whether the disrupted chromatin ensemble can return to a WT-like ensemble on biologically relevant interphase timescales, without becoming kinetically trapped in an aged or disorganized architectural state. Because the same LAD–NE interaction term that was removed in the model is restored in the recovery simulation, the result motivates an experimentally direct test based on rescue of productive LAD–lamina attachment. Note that the transition times to and from the *lamina-depleted* state are on the order of minutes, negligible compared to the duration of the interphase, which can last up to 16 hours for fruit fly somatic cells [112, 96]. Although the correspondence to biological time used here[109] cannot be considered exact, it is nevertheless expected to be accurate to roughly an order of magnitude, implying that upon restoration of the WT LAD-NE attractions, the WT state of chromatin is restored relatively fast.

The central experimentally testable prediction is that restoration of functional LAD–lamina interactions may be sufficient, under otherwise permissive conditions, to drive recovery toward a young/WT-like 3D chromatin architecture after lamina-dependent disruption. Importantly, the predicted intervention is not merely increased lamin abundance, but restoration of productive LAD association with the nuclear envelope. This distinction is essential experimentally: lamin expression and nuclear-envelope localization should be verified, but successful rescue of the model parameter should be assessed directly by LAD–lamina readouts such as DamID and/or fluorescence in situ hybridization (FISH)-based measurements of LAD radial positioning. If the model is correct, inducible lamin depletion followed by inducible lamin restoration should produce a reversible architectural trajectory: increased Hi-C map entropy and weakened chromosome territories after depletion, followed by a return to WT values after LAD–lamina contacts are restored. One practical route to test this prediction is as follows. The restoration of a youthful state of chromatin can be achieved by using GAL80(ts) or GeneSwitch-GAL4 inducible systems [7, 82] to express lamin in fruit fly cells. Lamin repression or depletion can be performed using the UAS-RNAi-lamin construct, while restoration of lamin expression can be accomplished using the UAS-lamin construct. To ensure robustness, lamin expression should be manipulated using both the RU486-inducible GeneSwitch-GAL4 and the temperature-sensitive GAL80(ts) systems. The key experimental requirement is to verify that the intervention restores LAD–lamina association, not only lamin protein abundance. If such experiments are performed, it will be important to determine how LADs reorganize spatially in response to these perturbations using FISH or DamID [111, 110]. In parallel, Hi-C data must be obtained to determine whether chromatin contact maps, Hi-C map entropy and chromosome-territory organization return toward their WT values after lamin restoration. Available experimental evidence is encouraging: lamin A re-expression was recently found to partially restore chromosome positioning, compaction, long-range interactions, and transcriptional profiles after lamin depletion in human stem cells[116].

Finally, if restoration of young-like 3D genome architecture contributes to cellular or organismal rejuvenation, these simulations motivate testing of whether restoration of functional LAD–lamina interactions in lamina depleted or aged flies improves aging-associated phenotypes, including Drosophila lifespan. Experimental studies have revealed a direct link between the function of lamins and organismal aging. *Drosophila* LamC mutations modeled after human progeroid LMNA mutations result in premature aging of adult flight muscles, including decreased levels of specific mitochondrial RNA transcripts and progressive mitochondrial degradation [61]. Knock-down of Dm0 in neurons led to a shorter lifespan, progressive impairment, and a loss of dopaminergic neurons in the protocerebral anterior medial cluster of the *D. melanogaster* brain [86]. The hypothesis that lamin restoration may improve lifespan or other aging-associated phenotypes could be tested by restoring WT lamin to physiologically normal levels, while also verifying recovery of functional LAD–lamina attachment. To be more precise, our model predicts that restoring functional LAD–nuclear envelope attachments — not merely lamin abundance — is sufficient to recover a WT-like 3D chromatin ensemble. In the case of rejuvenation of aged fruit fly, resetting the “young” epigenetic states of the LADs may be necessary as well to make LADs functional. For example, the aged-to-young cells ratio of the H3K27ac signal, a key marker for active enhancers and promoters, was inversely correlated with the basal H3K27ac signals, which means that regions that were strongly acetylated in young cells tended to lose H3K27ac with age, and vice versa. The observed blurring of the originally sharp H3K27ac pattern in dividing mouse embryonic fibroblasts demonstrates that the aging process corrupts epigenetic information [118]. Upon experimental lamina disruption in the *Drosophila* S2 cell line, the LADs become more acetylated at histone H3 and less compact after cells complete mitotic divisions [110]. The relevant epigenetic marks accumulate mainly during the G1 phase, remain relatively stable during the same interphase, and are removed from chromatin during mitosis [73]. However, epigenetic marks may also deteriorate in non-dividing cells during aging. For example, one study has shown increased chromatin accessibility and the loss of the heterochromatin mark H3K9me3 in the non-dividing excitatory neurons in the old mice compared to young ones [121]. In *Drosophila*, both H3K4me3 and H3K36me3 globally decrease across all actively expressed genes in non-dividing aging photoreceptors [44]. The senescence of *Drosophila* postmitotic fat body cells involves a reduction in Lamin B associated with age, which contributes to the loss of heterochromatin and the derepression of genes involved in immune responses [15]. Therefore, if some components of epigenetic aging accumulate during interphase, then partial architectural rejuvenation may be experimentally achievable in non-dividing cells. This highlights a knowledge gap in the direct mapping of epigenetic changes within LADs during normal aging in non-dividing cells, such as neurons. Studies of neuronal aging show lamina weakening, compartment shifts, and heterochromatin erosion [55, 120], but the analysis of histone and DNA methylation has not yet been conducted.

#### 3.2.2 Recovery from a limiting case “*lamina-depleted + compartment-disrupted*” state

Encouraged by the model prediction that lamina-depletion-induced architectural changes can be largely reversed by restoring WT-like LAD–NE interactions, we next test the recoverability of the chromatin architecture in an extreme limiting case. This stress test is designed to explore the extent of disorganization of a 3D genome architecture that can still be reversed on timescales much shorter than the duration of interphase. To this end, we consider a hypothetical fruit fly nucleus in which both all LAD–NE attractive interactions and all TAD–TAD attractive interactions have been switched off, see Methods. The resulting state of fruit fly chromatin, which we dub “*lamina-depleted + compartment-disrupted*” for notational simplicity, is represented by four separate self-avoiding polymer chains under confinement (the two chromosomes 2 and 3, plus the two chains corresponding to chromosome “X” and “4” confined within the NE).^b^

As before, we begin our simulation with 3 hrs in the WT state, Fig. 6, followed by an instantaneous switch off of all biologically relevant, and carefully tuned [109] attractive interactions within the entire chromatin at the TAD resolution, which yields a strongly simplified *lamina-depleted + compartment-disrupted* state dominated by polymer connectivity, confinement, and excluded-volume effects. Apart from near-diagonal contacts arising from chain connectivity, the contact map loses much of the WT-specific long-range structure, such as compartmentalization; see the upper triangle of the middle Hi-C map inset in Fig. 6. It is therefore particularly noteworthy that the original WT state of the chromatin is essentially restored within minutes upon restoring the relevant interactions within the chromatin and with the NE, as judged by contact-map entropy, Hi-C map similarity, and chromosome-territory intermingling. As in the *WT* → *lamina-depleted* → *rejuvenated* transition discussed above, the state recovered form *lamina-depleted + compartment-disrupted* is nearly indistinguishable from the original WT state by bulk Hi-C contact-map similarity. Qualitatively, the Hi-C maps of the two states are visually indistinguishable at the resolution of the model. Quantitatively, the contact probability difference map (not shown) between the two consists of matrix elements of the order of 10^*−*4^ or smaller, that is 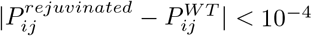. The Pearson correlation coefficient between the Hi-C maps of the original WT and the “rejuvenated after *lamina-depleted + compartment-disrupted*” states is 0.999987. Consistent with near-restoration of the WT-like Hi-C map, chromosome intermingling is reduced to approximately the WT level, Fig. 7, reversing the *lamina-depleted + compartment-disrupted*-induced increase in territory overlap. The subtle entropy ordering between *lamina-depleted* and *lamina-depleted + compartment-disrupted* states is discussed in the SI. A major increase and subsequent restoration to the WT value of cell-to-cell conformational heterogeneity along the trajectory shown in Fig. 6 are also discussed in the SI.

**Figure 6:**
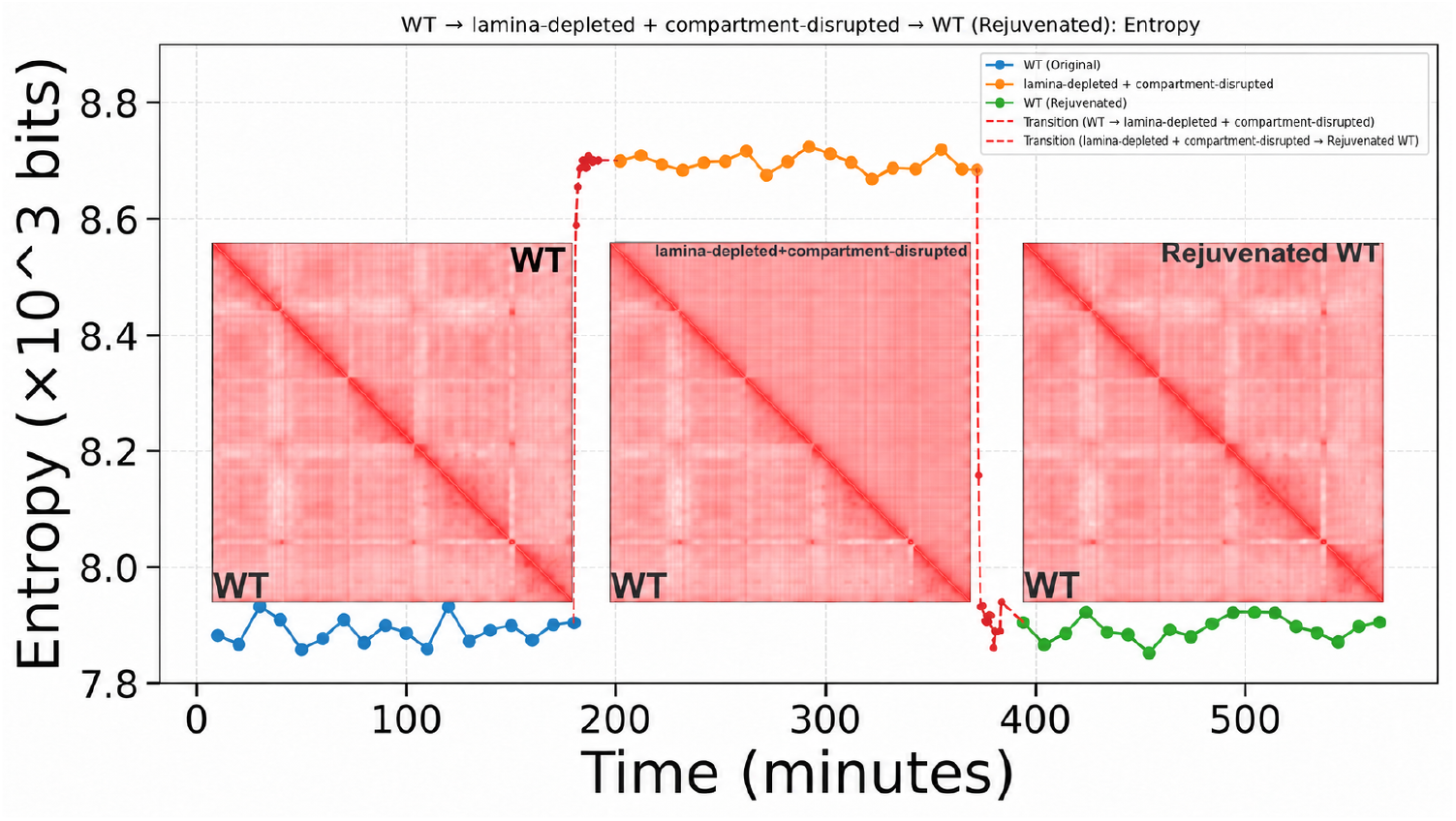
Transition from the WT to “*lamina-depleted + compartment-disrupted*” state of fruit fly is followed by its “rejuvenation” that fully restores the attractive interactions between all the elements of chromatin, including TADs, as well as between LADs and the nuclear envelope (NE), appropriate for the WT state and absent from the “*lamina-depleted + compartment-disrupted*” state. The transition to/from the “*lamina-depleted + compartment-disrupted*” state is initiated by an instantaneous switching off/on of all of the attractive interactions, see Methods. The corresponding Shannon entropy values, averaged over time, are ⟨*S*_WT_⟩ = 7890±5 bits, ⟨*Slamina-depleted + compartment-disrupted* ⟩ = 8699 ± 4 bits, and ⟨*S*_Rejuvenated WT_⟩ = 7895 ± 5 bits, where the error bars represent the standard error of the mean. Each of the three Hi-C maps shown as the insets is a bulk Hi-C map averaged over the full 3-hour window corresponding to each of the three states: WT, “*lamina-depleted + compartment-disrupted* “, or rejuvenated WT. To facilitate visual comparison, the lower triangle block of each matrix is the same original WT, while the upper triangle shows the state corresponding to the time interval.

**Figure 7:**
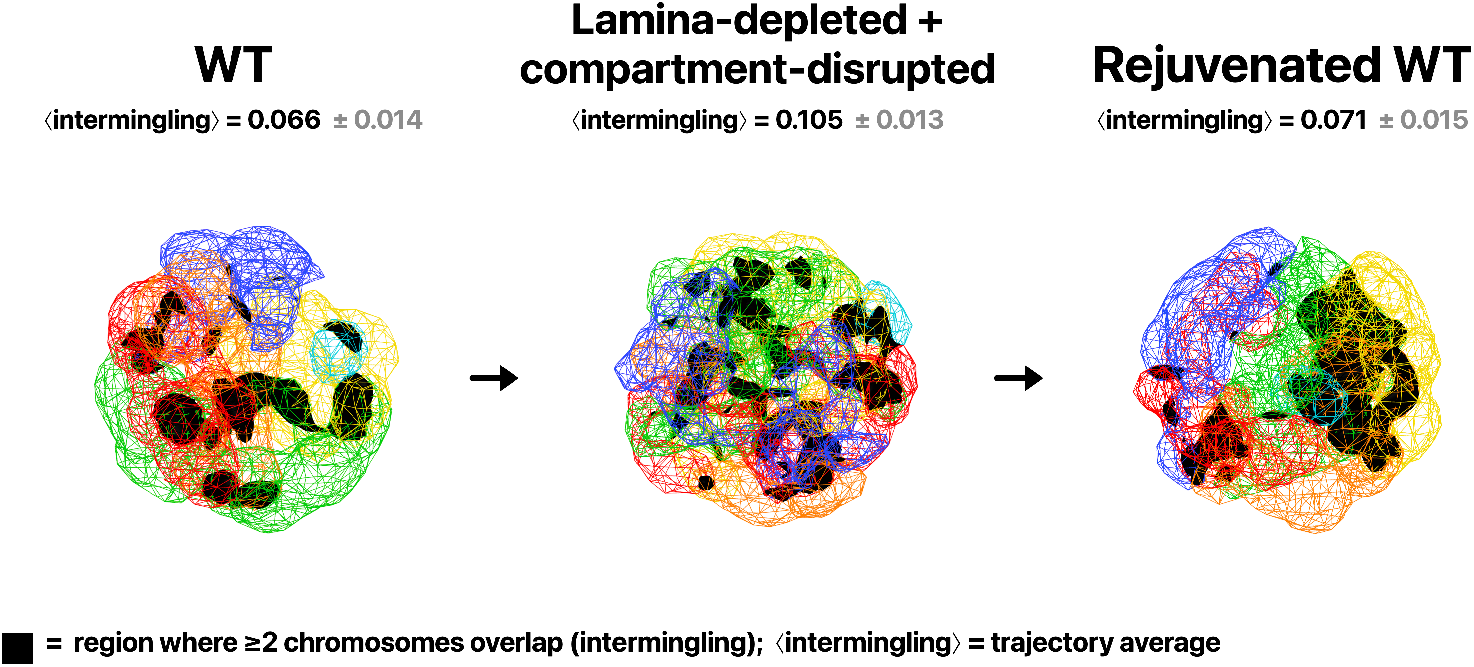
Chromosome territories along the WT → *lamina-depleted + compartment-disrupted* → WT rejuvenation pathway. The intermingling between chromosomes increases following transition to the *lamina-depleted + compartment-disrupted* state, and returns to its WT value upon the restoration of the attractive interactions between the TADs, and between LADs and the nuclear envelope (NE), present in the WT state and absent from the *lamina-depleted + compartment-disrupted* state. The snapshots shown are intended to illustrate each of the three states in Fig. 6; see also SI movies that show these states in dynamics. Each snapshot is selected so that its chromosome intermingling value closely approximates the trajectory average (shown above the snapshot) of the corresponding state. The intermingling regions are colored black. Each chromosome arm is colored differently, the gray sphere is the nucleolus. For this illustration, only one out of four experimentally observed mutual arrangements (topologies) of the chromosomes (CYSX-6S nucleus topology[109]) is shown. The errors are estimated as standard error of the mean.

The above results indicate that the WT-like 3D architecture can be recovered not only from the *lamina-depleted* state, but also from a more severely perturbed model state, *lamina-depleted + compartment-disrupted*, in which all attractive interactions have been switched off.

Within the computational model, the “rejuvenation” is achieved by switching back on the inter-loci (TAD-TAD) and chromatin-lamina interactions specific to the young nucleus. Can this prediction be tested experimentally? In fruit flies and mosquitoes, evolutionary conserved chromatin interactions can occur between genomic loci that are separated by several megabases [79, 72]. It was shown that CTCF (CCCTC-binding factor) and GAF (GAGA-associated factor), play direct roles in higher-order meta-domain interactions in *Drosophila* [79]. Further studies demonstrated that some specific long-range chromatin interactions depend on CTCF or transcription factors GAF and Vostok [79, 81, 42]. Manipulating these factors in fruit fly would affect the specific long-range chromatin interactions. Disruption of TADs would also disrupt TAD-TAD interactions. Therefore, in order to abolish much of the attractive TAD–TAD interaction landscape across the genome, the core architectural components of TADs should be the target in the experiment. In *Drosophila*, TADs arise from chromatin-state partitioning, active promoters/transcription, diverse architectural proteins and by boundary pairing [115, 8, 75]. In mammals, TADs are formed by cohesin-mediated loop extrusion at CTCF-defined boundaries [3, 95]. It has been shown that loss of cohesin eliminates most TADs and cohesin/CTCF loops and relaxes domain architecture in mammalian cells [3, 95]. This flattens intra-TAD structure globally and reduces structured TAD–TAD interaction patterns, although some compartmental or Polycomb-driven contacts persist [95]. Experiments on global disruption and restoration of TAD–TAD interactions in combination with manipulation lamina proteins have not yet been performed. If cells with broadly disrupted architectural interactions remain viable, future experiments could in principle test whether partial restoration of selected inter-loci and chromatin–lamina interactions promotes architectural recovery, although such experiments would be technically demanding.

## 4 Conclusions

The main motivation for this work was to explore the extent to which disorganization of chromatin conformation — a potentially important component of age-related deterioration of the epigenetic landscape — can be reversed by biologically meaningful manipulations, on timescales much shorter than interphase. Given the extraordinary complexity of hierarchical chromatin compaction in the interphase nucleus, our initial expectation was that even a partial restoration would be difficult.

The central conclusion and main testable prediction of this work is that the disorganization of the spatial state of fruit fly chromatin induced by lamina depletion, which mimics age-related changes, can be reversed to near-WT values, by the measured architectural metrics, when WT-like LAD–nuclear-envelope interactions are restored.

The recovery observed here should not be interpreted as a trivial consequence of restoring the same LAD–NE interaction term that was removed during lamina depletion. If interphase chromatin were an equilibrated polymer ensemble, then restoring the WT interaction landscape would be expected to eventually restore the WT equilibrium distribution, even if the recovery time-scales might be very long. However, large-scale interphase chromosome organization is widely understood to be far from equilibrium on cell-cycle timescales. In this context, the nontrivial result is that restoring WT-like LAD–NE interactions returns the disrupted chromatin ensemble to a WT-like architectural state on interphase timescales, without imposing individual TAD positions or chromosome territories. The result is robust to an even more drastic disruption of the 3D chromatin architecture: when both all LAD–NE attractive interactions and all TAD–TAD attractive interactions are switched off, subsequent restoration of the WT-like interactions also leads to recovery of the original WT 3D architecture of the chromatin.

The most direct biological prediction of this work is that restoration of functional LAD– lamina interactions, rather than lamin abundance alone, should promote recovery toward a young/WT-like 3D chromatin architecture after lamina-dependent disruption. This prediction can be tested by inducible lamina depletion and restoration in *Drosophila*, with DamID or FISH used to verify recovery of LAD–lamina association and Hi-C, chromosome-territory analysis, and single-cell Hi-C used to quantify architectural rescue. A concrete and realistic path to experimental verification of the prediction is proposed. Because lamina dysfunction is associated with aging and progeroid phenotypes, the result may contribute to fundamental understanding of the aging process, including how LAD–lamina interactions contribute to aging-associated 3D genome disorganization.

Thus, the youthful 3D genome architecture may be encoded not in a unique conformation, but in a distributed interaction landscape, including LAD–lamina contacts, TAD–TAD affinities, nuclear confinement, chromosome topology, etc. If aging degrades some of these organizing interactions without erasing the underlying genomic addresses, then restoring the interactions may allow chromatin to self-organize back into a youthful ensemble. Given the connection between lamin dysfunction and aging-associated phenotypes, we suggest that restoring functional LAD–lamina interactions may restore young-like 3D genome architecture and may be worth testing for effects on organismal aging phenotypes in fruit fly.

A methodological advance that facilitated our work is the notion of a Shannon entropy for Hi-C maps that we have introduced and rigorously developed here. The new quantity is introduced as a simple measure of the organization/disorganization of chromatin conformational states as defined by their Hi-C maps, and is designed to preserve the key features of entropy such as additivity for non-interacting systems. Throughout the work, we have used the Hi-C entropy as a simple and convenient metric of conformational disorder induced by age-associated perturbations such as lamina depletion. We have demonstrated that the new metric is sensitive to even subtle relevant changes in the Hi-C map, *e*.*g*., it can robustly distinguish between experimental WT and *lamina-depleted* Hi-C maps of *Drosophila melanogaster*, even though the visual differences are barely discernible. Importantly, the numerical differences in Hi-C entropy values upon lamina depletion predicted in our simulations of the chromatin evolution upon lamina depletion agree well with those estimated from the corresponding experimental Hi-C maps, supporting the model’s use in this context. Further testing in mammalian context, albeit very limited, is encouraging: the proposed Hi-C entropy is higher in the Progeria example analyzed here and tends to increase in selected chromosomes from human and mouse cerebellar cells across the lifespan.

The work has several limitations. First, near-restoration of selected 3D chromatin-architecture metrics has so far been demonstrated only in simulations of a female fruit fly nucleus, which has a relatively simple composition of only 4 chromosome pairs. Although the fruit fly is a higher eukaryote whose genome organization is complex, it is less complex than that of mammals with considerably more chromosomes. When our key conclusions are tested in mammals, this additional complexity of the 3D genome organization may result in the type of conformational rejuvenation described here being less complete and/or slower. A related methodological limitation is that models of chromatin considered here are at the TAD (~100 kb) resolution, which limits the amount of structural insight into the changes of the 3D chromatin architecture explored here. At the same time, the resulting computational efficiency of the simulation allowed us to simulate a biological ensemble of fruit fly nuclei on time scales of the interphase. The choice of TAD level resolution is justified by recent experiments that showed that lamina depletion affects only high-level chromatin organization, while the TAD structure remains intact[116]. Another limitation of the work is that we have only investigated interphase chromatin, that is, no passage through full cell cycle was considered. This limitation partly stems from the computational model used[109], which simulates only interphase chromatin, and partly from the limited availability of high-quality Hi-C maps covering entire lifespans. Our focus on the interphase is motivated in the Introduction, where we have argued that epigenetic rejuvenation approaches can benefit from avoiding DNA replication and the associated errors. Finally, we stress that despite encouraging results, we are not proposing Hi-C entropy as a *bona fide* epigenetic clock. Epigenetic clocks are complex, often relying on multiple features: further exploration is needed to determine whether the notion of Hi-C map entropy alone, or in conjunction with other related metrics[45, 66, 14], can be a useful addition to the existing arsenal of such features.

## Acknowledgment

The authors thank Dr. Longzhi Tan for kindly providing experimental life-spanning Hi-C maps of human and mouse cerebellar cells; Fatemeh Ghafouri for the help with making some of the pictures and movies illustrating various chromatin states. The work was supported in part by the National Institutes of Health, grant R35GM161622 to A.V.O. This work was also supported by the CeZAP Interdisciplinary Team-Building Pilot Grant Program, award number 202604. The authors acknowledge Advanced Research Computing at Virginia Tech for providing computational resources and technical support.

## Footnotes

a In Ref.[16] hysteresis was reported during a gradual decrease and subsequent increase of heterochromatin–lamina attraction in a polymer model of a single human chromosome. This result is not inconsistent with the reversibility observed here: hysteresis implies path dependence and metastability, but not permanent irreversibility. In the model of Ref.[16], the chromosome remained in the desorbed state over an intermediate range of restored lamina attraction, but returned to the adsorbed state once the reverse-transition threshold was crossed. Here, on the timescale of ~ 3 hrs, Fig. 4, we have not observed signs of instability[16] of the recovered WT state, although we can not exclude the possibility that it might be present on timescales shorter than the temporal resolution (10 mins) of our computational experiment. Further quantitative comparison with Ref. [16] is difficult for several reasons, including difficulty in comparing the relevant timescales (dimensionless Brownian time was used in Ref.[16]), the fact that we have simulated the entire fruit fly nucleus, while a single human chromosome was modeled in Ref.[16], and that here the original WT LAD–nuclear-envelope attraction is restored fully and abruptly, while the interaction strength was varied gradually in Ref. Ref.[16]

b This limiting *lamina-depleted + compartment-disrupted* state is conceptually related to the “desorbed-extended” state described in simulations of a human chromosome interacting with a model nuclear lamina[16]. In that model, simultaneous weakening of heterochromatin–lamina and heterochromatin– heterochromatin interactions produced an extended chromosome organization associated with progeroid cells. It is worth noting that the major loss of structure seen in the Hi-C map upon the *WT → lamina-depleted + compartment-disrupted* transition in our simulation, Fig. 6, is qualitatively similar to that in the *WT → Progeria* transition, see Fig. 2 and the relevant figure in Ref.[16]. We refrain from further quantitative comparison with Ref.[16] because the *lamina-depleted + compartment-disrupted* state is more extreme (all attractive TAD–TAD, LAD–NE, and nucleolar interactions are set to zero) than the “desorbed-extended” state, and the simulations here are performed for a whole-nucleus *Drosophila* genome ensemble rather than for a single human chromosome.

