## Supplemental Information for "Reversing aging-like 3D genome disorganization in a *Drosophila* interphase model"

July 2026

#### Post-processing of Hi-C maps for entropy analysis

**Row-normalized probability matrix.** For all Hi-C datasets used in this work, the Shannon entropy is computed from the row-normalized contact probability matrix $P_{ij}$. Given a raw Hi-C matrix $A_{ij}$, we apply the following (sequential) steps to obtain $P_{ij}$:

1. **Chromosome-aware exclusion (masking) of near-diagonal contacts**. Each locus is assigned a *D. melanogaster* chromosome label (2L, 2R, 3L, 3R, 4, X). Near-diagonal contacts with $|i-j| \le K$ are excluded *within* the same chromosome block, where in all calculations we used $K = 1$, unless otherwise stated. These elements are set to zero before the normalization.

2. **Row normalization**. For every row $i$, the remaining nonzero elements are normalized as
$$P_{ij} = \frac{A_{ij}}{\sum_k A_{ik}},$$
with rows that contained no valid contacts assigned a normalization factor of 1. This procedure ensures additivity of entropy across chromosomal blocks and avoids biases from near-diagonal structural contacts.

3. **Shannon entropy computation**. The entropy associated with each row is
$$s_i = -\sum_j P_{ij} \log_2 P_{ij},$$

and the total entropy is calculated as

$$S = \sum_i s_i.$$

4. **Visualization**. Normalized matrix elements $P_{ij}$ are visualized on a logarithmic color scale using a white–red colormap, where masked elements ($P_{ij} = 0$) are displayed in white. This allows structural differences—particularly long-range interactions—to be easily distinguished.

Unless otherwise specified, the above processing pipeline is applied uniformly to all of the Hi-C matrices used in this work, including WT, Progeria, *lamina-depleted* , *lamina-depleted + compartment-disrupted* , and rejuvenated states, ensuring a consistent comparison of entropy across experimental and simulated systems. Unless otherwise stated, all of the Hi-C maps shown in this work are row-normalized probability matrices $P_{ij}$.

### Effect of excluding $K$ near-diagonal elements from the Hi-C entropy estimate.

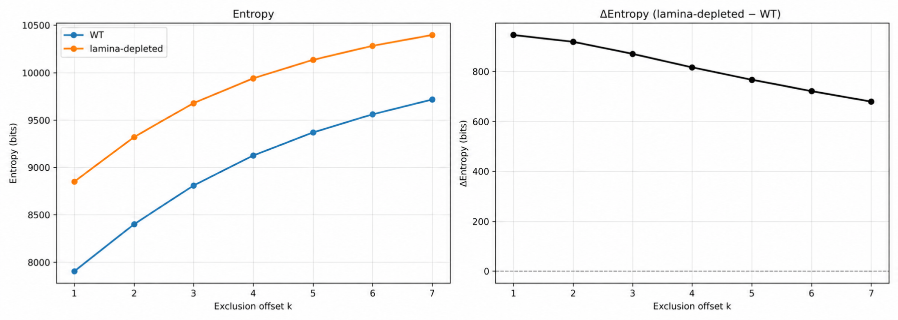


**Figure S1:** Behavior of Hi-C entropy $S(k)$ with increasing exclusion offset $k$ for WT and *lamina-depleted* chromatin contact maps. For each value of $k$, $S(k)$ was computed after excluding near-diagonal contacts within $|i - j| \leq k$, followed by row normalization of the remaining contact probabilities. **Left:** Entropy $S(k)$ for WT and *lamina-depleted* as a function of $k$. **Right:** Entropy difference $\Delta S(k) = S_{\textit{lamina-depleted}}\,(k) - S_{\mathrm{WT}}(k)$.

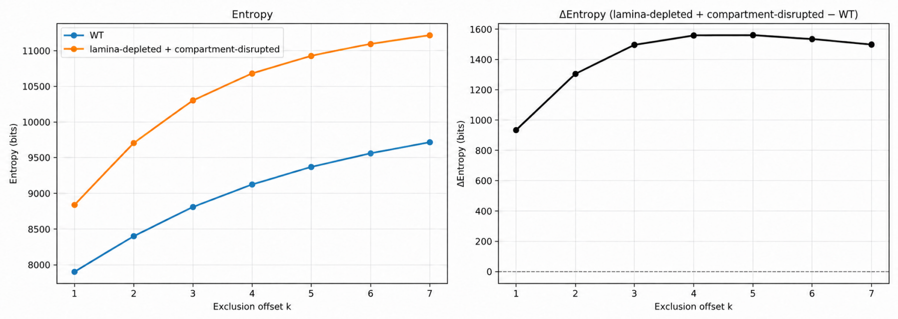


**Figure S2:** Behavior of Hi-C entropy $S(k)$ with increasing exclusion offset $k$ for the WT and *lamina-depleted + compartment-disrupted* chromatin contact maps. In each case, $S(k)$ is computed after excluding near-diagonal contacts within $|i - j| \leq k$, followed by row normalization of the remaining contact probabilities. **Left:** Entropy $S(k)$ for WT and *lamina-depleted + compartment-disrupted* as a function of $k$. **Right:** Entropy difference $\Delta S(k) = S_{\textit{lamina-depleted + compartment-disrupted}}\,(k) - S_{\mathrm{WT}}(k)$ as a function of $k$.

The dependence of $S(k)$ on the near-diagonal exclusion offset shows that transitions from the WT to the *lamina-depleted* or *lamina-depleted + compartment-disrupted* states alter different genomic-distance components of the WT Hi-C contact map. With the default $K = 1$, the *lamina-depleted* state has higher entropy than the *lamina-depleted + compartment-disrupted* state. Further analysis reveals that this behavior is dominated by near-diagonal contacts that remain included for $K = 1$: lamina depletion produces a more compact chromatin state and amplifies short-range contact contributions, while also causing stronger chromosome-territory intermingling. In contrast, transition to *lamina-*

*depleted + compartment-disrupted* removes the attractive TAD–TAD and LAD–NE interactions and visually erases much of the WT-specific long-range patterning, but does not produce the same near-diagonal contribution under the $K = 1$ normalization. When a broader near-diagonal band is excluded, for example $K = 4$, the entropy becomes more sensitive to medium- and long-range organization, and the entropy of the *lamina-depleted + compartment-disrupted* state becomes higher than that of the *lamina-depleted* state.

The ordering $S($*lamina-depleted*$) > S($*lamina-depleted + compartment-disrupted*$)$ seen at $K = 1$ is consistent with the chromosome-territory analysis. Although the *lamina-depleted + compartment-disrupted* Hi-C map visually loses more WT-specific long-range patterning, the *lamina-depleted* state shows stronger chromosome-territory intermingling. Thus, by a real-space measure of territorial disruption, lamina depletion is the more disruptive perturbation. This observation supports the interpretation that $K = 1$ Hi-C entropy is sensitive to a biologically meaningful component of architectural disorder, rather than simply failing to rank the visually least structured Hi-C map highest. At the same time, the reversal of the ordering at larger values of $K$ indicates that *lamina-depleted + compartment-disrupted* state more strongly disrupts medium- and long-range contact-map organization. Together, the entropy and intermingling analyses suggest that *lamina-depleted* and *lamina-depleted + compartment-disrupted* states represent somewhat different modes of chromatin disorganization rather than merely points on a single universal disorder scale. The latter conclusion is consistent with the finding from an earlier simulation study[2] that investigated the response of a single chromosome to altering chromatin-lamina interactions.

Importantly, the central conclusions of this work — entropy increases after disruption and returns toward WT after restoration of the WT interaction landscape — are robust to the choice of $K$.

### Entropy of the Hi-C map of a mammalian nucleus in the interphase.

For each of the three human chromosomes examined, Fig. S3, the Hi-C entropy increases monotonically across the sampled ages. The three corresponding mouse chromosomes did not show strictly monotonic behavior, although their entropy values tended to be higher at older ages – one can see this by averaging the entropy values over the last two vs. first two age points in Fig. S3.

These results provide additional support for the utility of the proposed definition of Hi-C map entropy $S(A_{ij})$ in studies that explore structural changes in chromatin with age. The results suggest that some age-associated changes in 3D genome architecture may be captured by contact-map disorder in Hi-C maps. In this context, the proposed Hi-C entropy may be a useful metric for aging-associated changes in 3D genome organization.

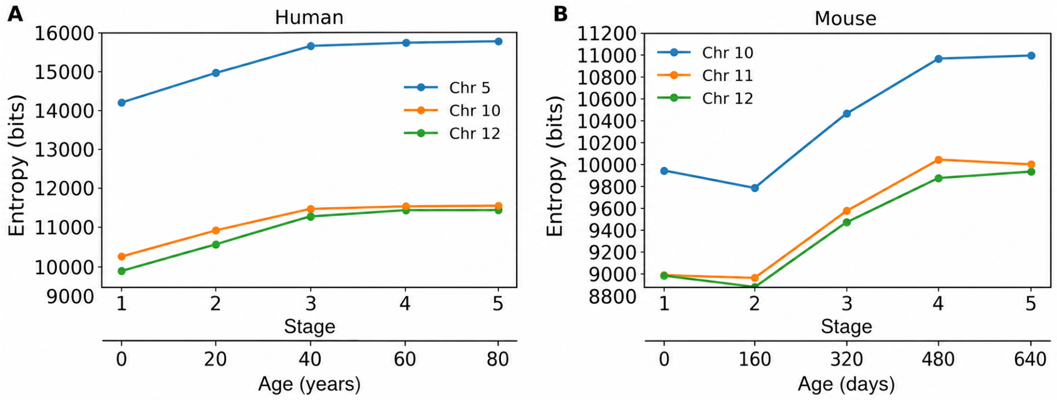


**Figure S3:** Hi-C entropy $S(t)$ across developmental stages for human (left) and mouse (right) chromosomes. Each panel shows Hi-C entropy of the three representative chromosomes from cerebellar granule cells selected and examined in Ref. [8]. The entropy is calculated as described in "Methods" using the original Hi-C data from Ref. [8], the maps are shown in Figs. S4 and S5. The stages, as defined in Ref.[8], are shown on the primary x-axis, while the secondary axis indicates the corresponding age of the organism (from 0 to the maximum investigated). The Hi-C entropy tends to increase with the age of the organism.

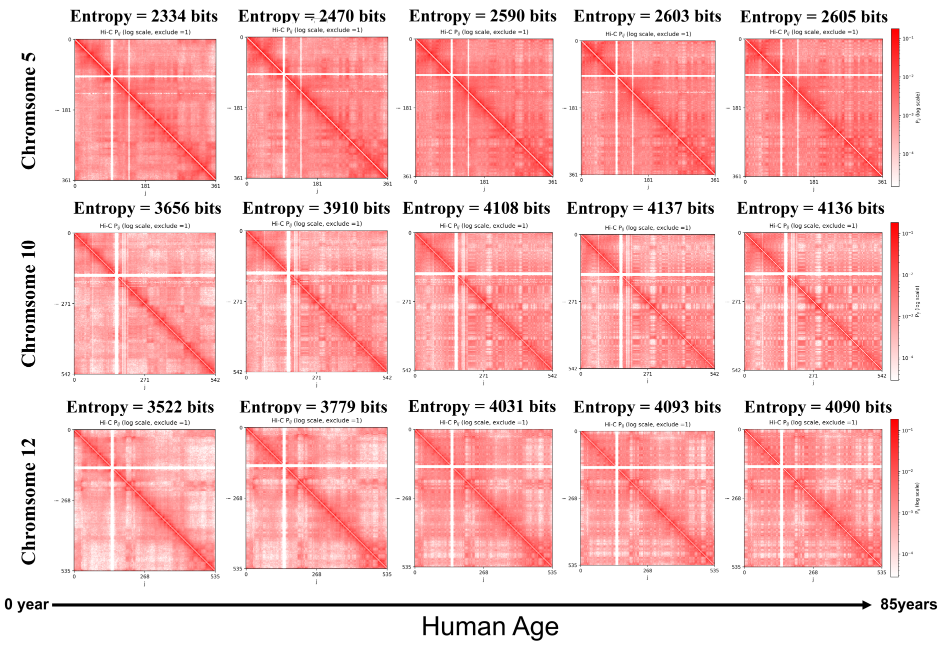


**Figure S4:** Entropy of Hi-C contact maps increases across sampled ages for selected human chromosomes shown here. Shown are contact probabilities corresponding to human chromosomes in the interphase (neurons: cerebellar granule cells) selected and examined in Ref. [8]. The original Hi-C data are from Ref. [8]. The entropy values are shown as insets.

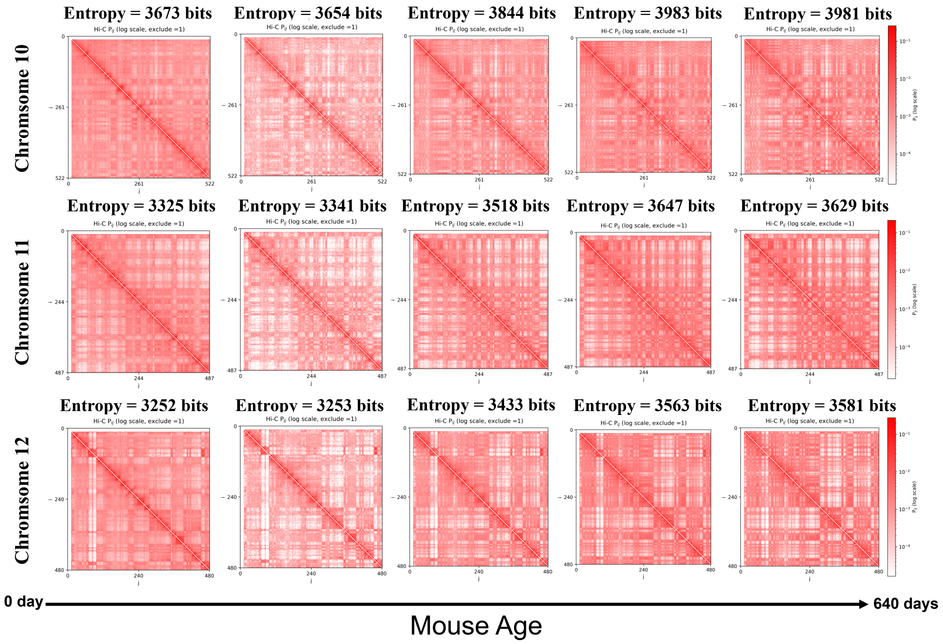


**Figure S5:** Entropy of Hi-C contact maps tends to increase with age for selected mouse chromosomes shown here. Shown are contact probabilities corresponding to mouse chromosomes in the interphase (neurons: cerebellar granule cells) selected and examined in Ref. [8]. The original Hi-C data are from Ref. [8]. The entropy values are shown as insets.

### Computing chromosome intermingling: temporal sampling and statistics.

For each simulation, the intermingling index $I$ is evaluated on 1000 equidistant snapshots sampled uniformly from the entire simulated trajectory corresponding to the given state of chromatin (frame indices round$[\,i\,(T-1)/999\,]$, $i = 0 \ldots 999$; $T = 109{,}183$ frames $\approx$ $10{,}800\,\mathrm{s} \approx 3\,\mathrm{h}$; stride $\approx 109$ frames $\approx 10.8\,\mathrm{s}$). The reported $\langle I \rangle$ is the mean of these 1000 per-frame values, and the quoted uncertainty is their population standard deviation,

$$\sigma_I \;=\; \sqrt{\frac{1}{N}\sum_{t=1}^{N}\big(I_t - \langle I \rangle\big)^2}, \qquad N = 1000, \tag{1}$$

### Visualization of chromosome territories and intermingling.

Territory renderings are produced from individual snapshots: each arm's bead coordinates are convolved with an isotropic Gaussian (width $0.18\,\sigma$) onto a $0.12\,\sigma$ grid, and a fixed iso-level contour of the resulting density, drawn as a colored wire frame mesh, defines each arm's visible territory; the overlap (intermingling) volume is rendered as a solid surface of the field given by the second-largest of the six per-arm densities at each voxel. Density fields, the intermingling index, and frame selection were implemented in custom Python 3 scripts. Three-dimensional rendering used VMD 2.0.0a7[3] with the Isosurface and Licorice representations, ray-traced with the bundled Tachyon renderer (12× anti-aliasing, ambient occlusion). The original simulation trajectories are read in VTF format. Scripts to produce the movies and to compute the intermingling index are available at `https://github.com/Onufriev-Lab/intermingling_movie_kit`

### Conformational Heterogeneity (C.H.) based on Hi-C contact probabilities

The bulk Hi-C entropy introduced in this work, can not, by itself, measure cell-to-cell conformational heterogeneity. To quantify cell-to-cell heterogeneity in chromatin 3D organization, we adopt the concept of Conformational Heterogeneity, *C.H.*, introduced in Ref. [5], where *C.H.* was originally defined as the standard deviation of average spatial Euclidean distances $\langle R_s^{(k)} \rangle$ between genomic loci separated by distance $s$ over an ensemble of single cells [5]:

$$C.H.(s) = \sqrt{\frac{1}{n} \sum_{k=1}^{n} \left( \langle R_s^{(k)} \rangle - \langle R_s \rangle \right)^2}, \tag{2}$$

where $\langle R_s^{(k)} \rangle$ is the average spatial distance between all pairs of loci at distance $s$ in the $k$-th conformation, and $\langle R_s \rangle$ is the average over all $n$ conformations (cells).

In this work, we replace the mean physical distance $\langle R_s^{(k)} \rangle$ with the mean row-normalized Hi-C contact probability $\langle P_s^{(k)} \rangle$ for each sample, so that the *C.H.* can be computed directly from experimental Hi-C probabilities. The revised formula reads:

$$C.H.(P_s) = \sqrt{\frac{1}{n} \sum_{k=1}^{n} \left( \langle P_s^{(k)} \rangle - \langle P_s \rangle \right)^2}, \tag{3}$$

where $\langle P_s^{(k)} \rangle$ is the mean probability of contact at genomic distance $s$ for the $k$-th sample (after row normalization and exclusion of near-diagonal contacts with $|i - j| \leq 1$), and $\langle P_s \rangle$ is the average value across all samples.

This formula yields a direct measure of *C.H.* based on the contact probabilities available from the Hi-C map, thus bypassing the need for a full 3D structure in Euclidean space, which can be difficult or impossible to obtain.

In this work, each of the five distinct states of chromatin: WT, *lamina-depleted + compartment-disrupted* , *lamina-depleted* , and both "rejuvenated" states is represented by sets of model nuclei, generated as described in the main text. Each set, which comprises multiple chromatin configurations, corresponds to one data point in Figs. 4 and 6 of the main text. The (average) Hi-C map corresponding to each set is used as a data point in Eq. 3. In the implementation of Eq. 3 used in this work, both upper and lower matrix triangles are utilized after normalization.

#### Rejuvenation reduces cell-to-cell conformational heterogeneity of chromatin.

It was previously hypothesized[9] that the removal of attractive LAD-NE interactions, which stabilize overall chromatin organization, can lead to increased heterogeneity of chromatin conformations. Here we explore whether the trend can also be seen in the more drastic case of the *lamina-depleted + compartment-disrupted* state, and whether it can be reversed by restoring the WT LAD-NE and TAD-TAD attractions. To this end, we employ a newly developed [5] quantitative metric of cell-to-cell *conformational heterogeneity*, *C.H.*, of chromatin organization in 3D. By construction, $C.H.(s)$ averages out conformational variability within each given nucleus, thereby focusing specifically on cell-to-cell variability. That is, $C.H.(s)$ would yield exactly zero for a hypothetical case of identical nuclei, even if the chromatin conformations within each nucleus were highly variable. Here, we adapt the metric so that it can be computed directly from the Hi-C map, which has multiple practical advantages, see above. Specifically, $C.H.$ at genomic separation $s$ is defined as the standard deviation of the average probability of contact between chromatin loci separated by genomic distance $s$, Eq. 3.

To summarize the overall heterogeneity by a single number per chromatin state, we average $C.H.(P_s)$ over all available genomic separations $s$ on the *whole* nucleus, Table 1. The average provides a compact, global measure of conformational variability that can be directly compared across nuclear states.

**Table 1: Whole-genome mean conformational heterogeneity (averaged over $s$) for each state.** Values are reported as unweighted mean ± jackknife standard error across replicate Hi-C maps ($n = 18$ per state) and are ordered from highest to lowest.

| **State** | $\overline{\mathrm{CH}}$ ($\times 10^{-5}$) |
|---|---|
| *lamina-depleted + compartment-disrupted* | $9.35 \pm 0.46$ |
| *lamina-depleted* | $6.78 \pm 0.24$ |
| Rejuvenated from *lamina-depleted* | $6.60 \pm 0.32$ |
| WT | $6.29 \pm 0.24$ |
| Rejuvenated from *lamina-depleted + compartment-disrupted* | $6.15 \pm 0.47$ |

Throughout the whole nucleus, conformational heterogeneity increases significantly after severe chromatin disruption (*lamina-depleted + compartment-disrupted* ), with a small uptick observed after the less drastic *lamina-depleted* perturbation. What is most important in the context of this work, is that restoration of the WT interactions reduces the average $C.H.(s)$ in all cases, returning the system toward the WT levels.

Chromosome-arm–resolved $C.H.(s)$ values, Fig. S6, show that, while the magnitude and detailed distance dependence of $C.H.(s)$ vary across individual chromosome arms, the over-all conclusions based are broadly consistent with the whole-genome summary based on Table 1. These arm-specific results further support the use of a global $C.H.$ measure in the main text, while highlighting heterogeneity in the contributions of different chromosomal regions.

Further analysis, Fig. S7, reveals that the most drastic changes in cell-to-cell conformational heterogeneity occur at the largest genomic distances, larger than 10 Mb, with the effect essentially disappearing for the smallest distances of less than 1 Mb. The rejuvenation effect – return to the WT values – is seen for both the largest and the intermediate genomic distances, though it is much less pronounced for the latter since the increase in $C.H.$ is itself not nearly as pronounced.

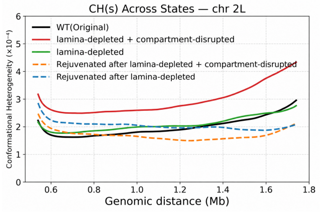


(a) chr 2L

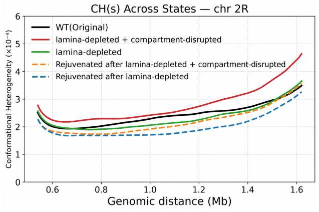


(b) chr 2R

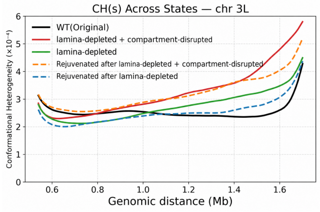


(c) chr 3L

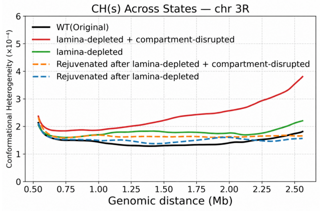


(d) chr 3R

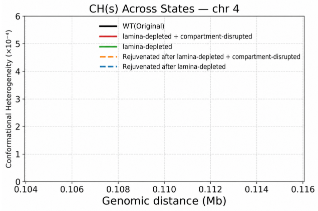


(e) chr 4

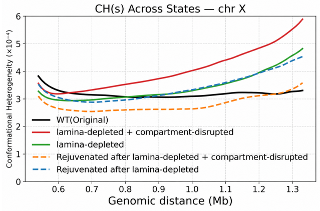


(f) chr X

**Figure S6: Chromosome-arm–resolved conformational heterogeneity (CH) across nuclear states.** CH(s) is shown as a function of genomic separation for individual chromosome arms (2L, 2R, 3L, 3R, and X). For each arm, the maximum genomic separation is limited by the arm length, and the curves terminate accordingly. No curves are shown for the very small chromosome 4 due to insufficient genomic extent to support a meaningful analysis.

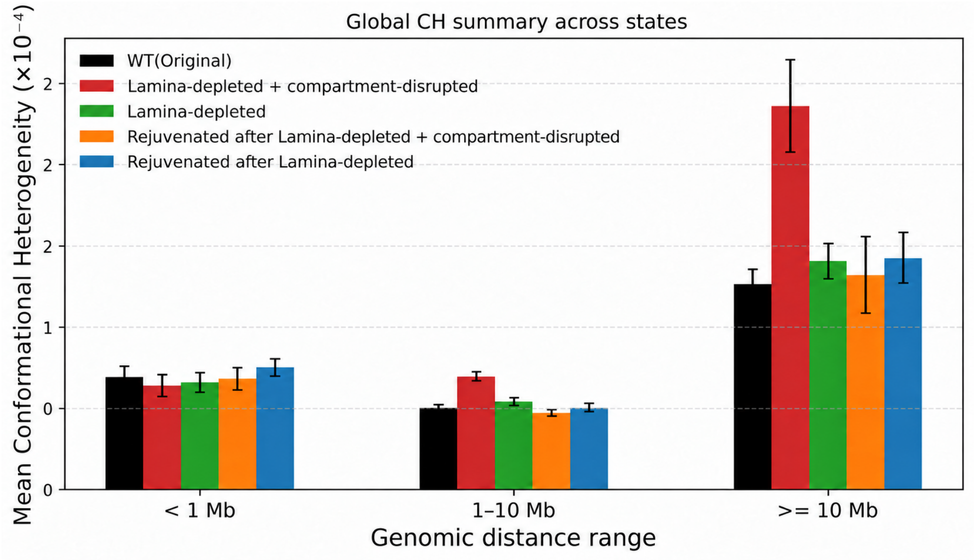


**Figure S7: Whole-genome comparison of conformational heterogeneity (C.H.) across nuclear states.** For each Hi-C map, the contact probability matrix is row-normalized after excluding the diagonal. *C.H.* is defined as the standard deviation of $P(s)$ across replicate maps for each state. The bars report *C.H.* averaged within three genomic distance ranges ($< 1$ Mb, $1$–$10$ Mb, and $\geq 10$ Mb). Error bars indicate jackknife standard errors across replicate Hi-C maps.

In summary, we suggest that cell-to-cell conformational heterogeneity of chromatin increases substantially when both chromatin–NE and TAD–TAD interactions are disrupted, while disruption of chromatin–NE interactions alone leads to only very modest increase. Thus, the substantial loss or dysfunction of lamins during aging or disease [1, 4, 6] may contribute to increased disorder in gene expression by enhancing variability in global chromatin architecture. These findings highlight a key role of chromatin–NE and TAD–TAD interactions in the establishment of non-random three-dimensional chromatin organization and in the preservation of its integrity against the inherent cell-to-cell variability of chromatin conformations. Most importantly, our simulations further suggest that the increase in conformational heterogeneity associated with disruption of chromatin–NE interactions, and especially in the extreme case of disruption of both chromatin–NE and TAD–TAD interactions, can be mostly reversed by restoring these interactions to WT values.

This prediction, which can be tested using single-cell Hi-C experiments or related single-cell 3D genome assays[10], contributes to our understanding of aging and may help explain, at least in part, why the epigenetic state of older differentiated cells is often less well defined than that of younger cells, a proposed hallmark of epigenetic aging[7]. Significant loss or dysfunction of lamins during aging or disease [1, 4, 6] can contribute to increased disorder in gene expression by enhancing variability in the global chromatin architecture. We suggest that the proposed association between cell-to-cell heterogeneity of 3D genome architecture and cell age is worth exploring.

[2] Michael Chiang, Davide Michieletto, Chris A Brackley, Nattaphong Rattanavirotkul, Hisham Mohammed, Davide Marenduzzo, and Tamir Chandra. Lamina and heterochromatin direct chromosome organisation in senescence and progeria. *bioRxiv*, page 468561, 2018.
