## Supplemental Movies for "Reversing aging-like 3D genome disorganization in a *Drosophila* interphase model": README.pdf

July 2026

### The dynamic model of the fruit fly nucleus.

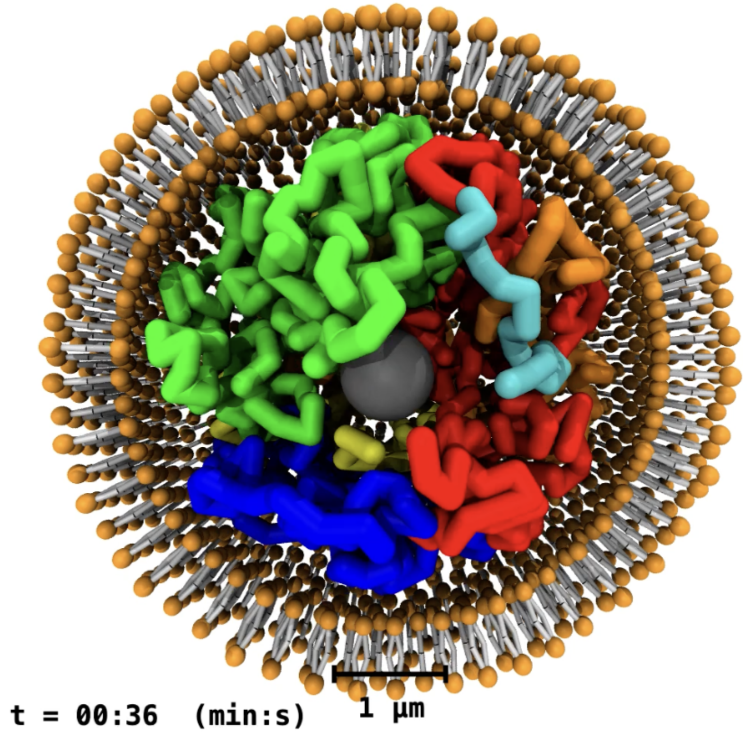


**Figure SMovie1:** A movie illustrating about 30 seconds of the computed temporal evolution of the entire fruit fly chromatin confined within the nucleus during interphase. Only one out of four experimentally observed mutual arrangements (CYSX-6S[1]) of the chromosomes is shown. Each chromosome arm is colored differently, the gray sphere is the nucleolus.

### Recovery from *lamina-depleted* state

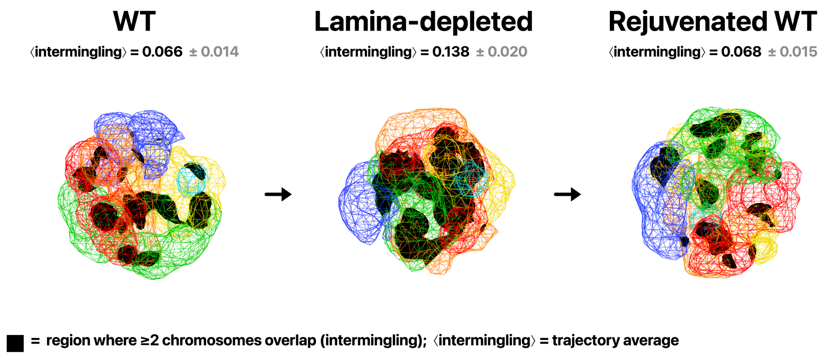


**Figure SMovie2:** Temporal evolution of chromosome territories along the WT → *lamina-depleted* → WT rejuvenation pathway (about 10 seconds per state). The intermingling between chromosomes increases following transition to the *lamina-depleted* state, and returns to its WT value upon the restoration of the attractive interactions between LADs and the nuclear envelope (NE), present in the WT state and absent from the *lamina-depleted* state. The intermingling regions are colored black. Each chromosome arm is colored differently, the gray sphere is the nucleolus. For clarity, only one out of four experimentally observed mutual arrangements (CYSX-6S[1]) of the chromosomes is shown.

### Recovery from " *lamina-depleted + compartment-disrupted* " state

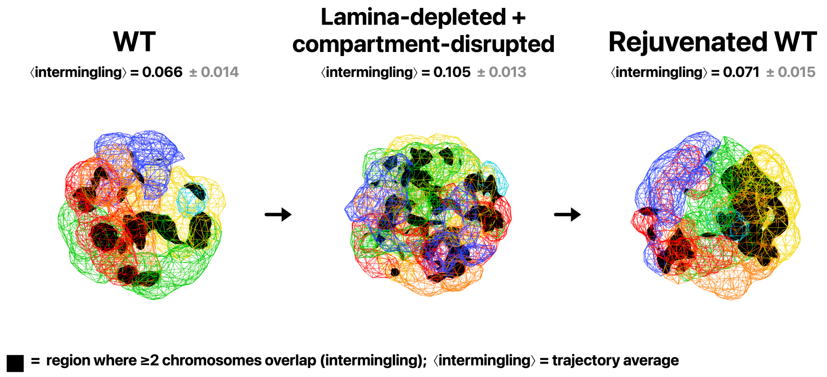


**Figure SMovie3:** Temporal evolution of chromosome territories along the WT → *lamina-depleted + compartment-disrupted* → WT rejuvenation pathway (about 10 seconds per state). The intermingling between chromosomes increases following transition to the *lamina-depleted + compartment-disrupted* state, and returns to its WT value upon the restoration of the attractive interactions between the TADs, and between LADs and the nuclear envelope (NE), present in the WT state and absent from the *lamina-depleted + compartment-disrupted* state. The intermingling regions are colored black. Each chromosome arm is colored differently, the gray sphere is the nucleolus. For clarity, only one out of four experimentally observed mutual arrangements (CYSX-6S[1]) of the chromosomes is shown.
